# Characterization of *Rose* rosette *virus* and development of reverse genetic system for studying virus accumulation and movement in whole plants

**DOI:** 10.1101/712000

**Authors:** Jeanmarie Verchot, Venura Herath, Cesar D. Urrutia, Mathieu Gayral, Kelsey Lyle, Madalyn K. Shires, Kevin Ong, David Byrne

**Author notes:** Address correspondence to Jeanmarie Verchot. Department of Plant Pathology & Microbiology, Texas A&M University, College Station, Texas, USA.

## Abstract

*Rose rosette virus* (RRV) is an *Emaravirus*, a negative-sense RNA virus with a 7-segmented genome that is enclosed by a double membrane. While the genome sequences of many emaraviruses are reported, there is negligible information concerning virus replication and movement in host plants. Computational methods determined that RNA1 encoded the RNA dependent RNA polymerase (RdRp), RNA2 encoded glycoprotein precursor, and the RNA3 encoded the nucleocapsid (N), all share significant homologies with similar proteins of the Orthobunyavirus family. The RRV terminal UTR sequences are complementary and share significant identity with the UTR sequences of Bunyamwera virus. We report a minireplicon system and a full length infectious clone of RRV, which are the first for any emaravirus species. The photoreversible fluorescent iLOV protein was used to replace the RNA5 open reading frame (R5-iLOV). We demonstrate that agro-infiltration of *Nicotiana benthamiana* leaves to deliver RNA1, RNA3, and R5-iLOV cDNAs led to iLOV expression. A mutation was introduced into the RdRp active site and iLOV expression was eliminated. Delivery of four segments or seven segments of the RRV infectious clone produced systemic infection in *N. benthamiana* and rose plants. iLOV was also fused to the glycoprotein precursor (R2-iLOV). Using confocal microscopy, the R2-iLOV was seen in spherical bodies along membrane strands inside *N. benthamiana* epidermal cells. This new technology will enable future research to functionally characterize the RRV proteins, to study the virus-host interactions governing local and systemic infection, and examine the subcellular functions of the Gc.

**IMPORTANCE:** RRV has emerged as a severe threat to cultivated roses, causing millions of dollars in losses to commercial producers. The majority of the viral gene products have not been researched or characterized until now. We constructed a minireplicon system and an infectious clone of the seven-segmented RRV genome that is contained in a binary vector and delivered by Agrobacterium. This technology has been slow to develop for viruses with negative-strand RNA genomes. It has been especially tricky for plant viruses with multicomponent negative-strand RNA genomes. We report the first reverse genetic system for a member of the genus *Emaravirus, Rose rosette virus* (RRV). We introduced the iLOV fluorescent protein as a fusion to the Gc protein and as a replacement for the open reading frame in genome segment 5. This game-changing reverse genetic system creates new opportunities for studying negative-strand RNA viruses in plants.

## INTRODUCTION

Among the most significant technological advances for research into virus life cycles has been the development of cDNA copies of viral genomes that can be reverse transcribed to produce infectious virus (1). This technology enables the genetic manipulation of cDNA for reverse genetic analysis of the virus life cycle, and investigations into the molecular basis of virus-host interactions for susceptibility and immunity. The infectious cDNA technology is routinely used to study positive-strand RNA viruses. Infectious virus cDNAs are inserted into plasmids by fusing the exact 5’ end of the virus genomic cDNA to the bacteriophage T7, SP6, RNA pol I, or RNA pol II promoters (2–5). A transcriptional terminator or hepatitis delta virus ribozyme (HDRz) can be located at the 3’ ends to produce transcripts with exact 3’ ends (6–8). The second advance in infectious clone technology was the discovery that genetic sequences encoding fluorescent proteins can be introduced into specific locations within the viral genome to visualize the effects of mutations on virus replication in reverse genetic experiments (9–16). Among plant RNA viruses, green fluorescent protein (GFP) and derivatives, or iLOV genes, have been incorporated into viral genomes (17). Combining visual markers of infection with reverse genetic technology has been a powerful combination for functional imaging to uncover critical roles of viral proteins in the virus life cycle. These combined technologies have been central to the discovery of molecular plant-virus interactions occurring in susceptible hosts or that govern gene-for-gene resistance.

Infectious clone technology has been tricky to develop for plant viruses with negative-strand RNA genomes because the naked genomic or antigenomic RNAs are not able to initiate infection by themselves. The minimum infectious unit for viruses with negative-strand RNA genomes requires a ribonucleoprotein (RNP) complex composed of viral genomic RNA and RNA dependent RNA polymerase (RdRp). For plant infecting viruses, a movement protein (MP) is also essential for whole plant infection. Considering the many successful examples of infectious clones developed for negative-strand RNA viruses such as species belonging to the families *Rhabdoviridae, Paramyxoviridae, and Filoviridae* (18–20), infection is achievable by delivering viral genomic cDNAs that produce transcripts functioning as the antigenomic RNAs (agRNA). Plasmids encoding the nucleocapsid core or subunits of the viral polymerase are co-delivered with the agRNA encoding cDNAs that successfully spur the replication process. The first infectious clone of a plant-infecting negative-strand RNA virus was *Sonchus yellow net virus* (SYNV) (18, 19, 21). This construction consists of a binary vector containing the SYNV full-length cDNA fused to a duplicated Cauliflower mosaic virus (CaMV) 35S promoter. Additional binary plasmids expressing the NC (nucleocapsid), P (phosphoprotein), and L (polymerase) proteins are co-delivered to leaves for active SYNV infection.

The order *Bunyavirales* includes nine families of enveloped viruses with segmented negative-strand RNA genomes. These viruses infect animals, plants, and invertebrates. Four families contain viruses that cause disease in humans, including *La Cross virus* (LACV), *Rift valley fever virus* (RVFV), *Crimean-Congo hemorrhagic fever virus* (CCHFV) and *Hantaan virus* (HTNV) (22, 23). Three families within this order encode viruses that infect plants: *Fimoviridae, Phenuiviridae*, and *Tospoviridae*. Their genomes consist of three to eight segments with the largest segment (named L or RNA1) encoding the RNA dependent RNA polymerase (RdRp). For viruses with three genome segments, the M segment encodes two glycoproteins (Gn and Gc), and the S segment encodes the nucleocapsid (named NC or N for different genera or species) protein. The untranslated regions (UTRs) at the 3’ and 5’ ends direct transcription of mRNA and replication to generate agRNA as intermediates for synthesis of further genomic RNAs. Full-length infectious clones and “minireplicon” systems are available for studying the bunyavirus replication and transcription at the cellular level (24). These systems are successful for achieving RNA synthesis when the RdRp and NC are expressed from cDNA plasmids as the minimum requirements to achieve transcription and replication-competent RNP particles. Co-transfecting plasmids encoding the viral glycoprotein precursor stimulates the packaging of the RNPs into virus-like particles.

*Rose rosette virus* (RRV) is a negative-strand RNA virus with seven genome segments and belongs to the *Emaravirus* genus within the family *Fimoviridae*. Beyond reporting the genome sequence for RRV, there is very little known about the functional roles of proteins encoded by each RNA segment. Roses are the economically most important ornamental plants in the *Rosaceae* family. RRV has been devastating roses and the rose industry in the USA, causing millions of dollars in losses (25). Typical symptoms of RRV include rapid stem elongation, breaking of axillary buds, leaflet deformation and wrinkling, bright red pigmentation, phyllody, and hyper-thorniness (30,31). Currently, researchers rely on viruliferous eriophyid mites to deliver RRV to plants, and the mechanical delivery of RRV to test plants has not consistently effective. Here we describe the molecular properties of RRV and report a minireplicon system that can be used to study the viral components of the replication cycle. We also report an infectious clone of RRV that can be used to study virus-host interactions in whole plants. Each cDNA segment is fused to the duplicated CaMV 35S promoter and a 3’ HDRz. We successfully introduced the iLOV gene in two viral genome locations to study virus replication and glycoprotein functions in plant cells. We demonstrate the utility of this enhanced visual reporter system for reverse genetic studies of infection and for visualizing the virus replication cycle in *N. benthamiana* and rose plants.

## RESULTS

### Rose rosette virus RNA-dependent RNA polymerase share similarities with orthobunyaviruses

For most *Bunyavirales* members, the RdRp and N proteins constitute the minimal protein machinery for genome replication and transcription. The terminal UTRs are also critical for replication and transcription. The 5’ and 3’ UTRs are complementary and form a “panhandle” required for efficient protein synthesis in mini-replicon systems (26). Thus, identifying the RdRp and N proteins and characterizing the UTR regions was essential for building an RRV mini-replicon system as well as an infectious cDNA clone.

To characterize the viral RdRp, we also compiled the RdRp sequences and accessions of classified bunyavirus species provided in the 2019 ICTV taxonomic update (Tables S1)(22). We performed sequence analysis and alignments using Muscle v 3.8.425 within the Geneious Prime® 2019.2.1 software. Fig. 1 presents a phylogenetic tree of the RdRp sequences from 70 species that was constructed using W-IQ-Tree with 1000 ultrafast bootstraps (28). Members of the *Emaravirus* genus (Family *Fimoviridae*) clustered with the *Orthobunyavirus* genus (Family *Peribunayviridae*), represented by LACV and *Bunyamwera* (BUNV) virus in a common clade. The closest neighboring cluster were species within the genus *Hantavirus* and *Orthophasmavirus* (Families *Hantaviridae* and *Phasmaviridae*, respectively). The two genera of plant infecting viruses, named *Orthotospovirus (*Family *Tospoviridae)* and *Tenuivirus* (Family *Phenuiviridae*), were distant from the *Emaravirus* and clustered with members of the *Phlebovirus* genus (Family *Phenuiviridae*) such as RVFV (Fig. 1).

**Fig. 1.**
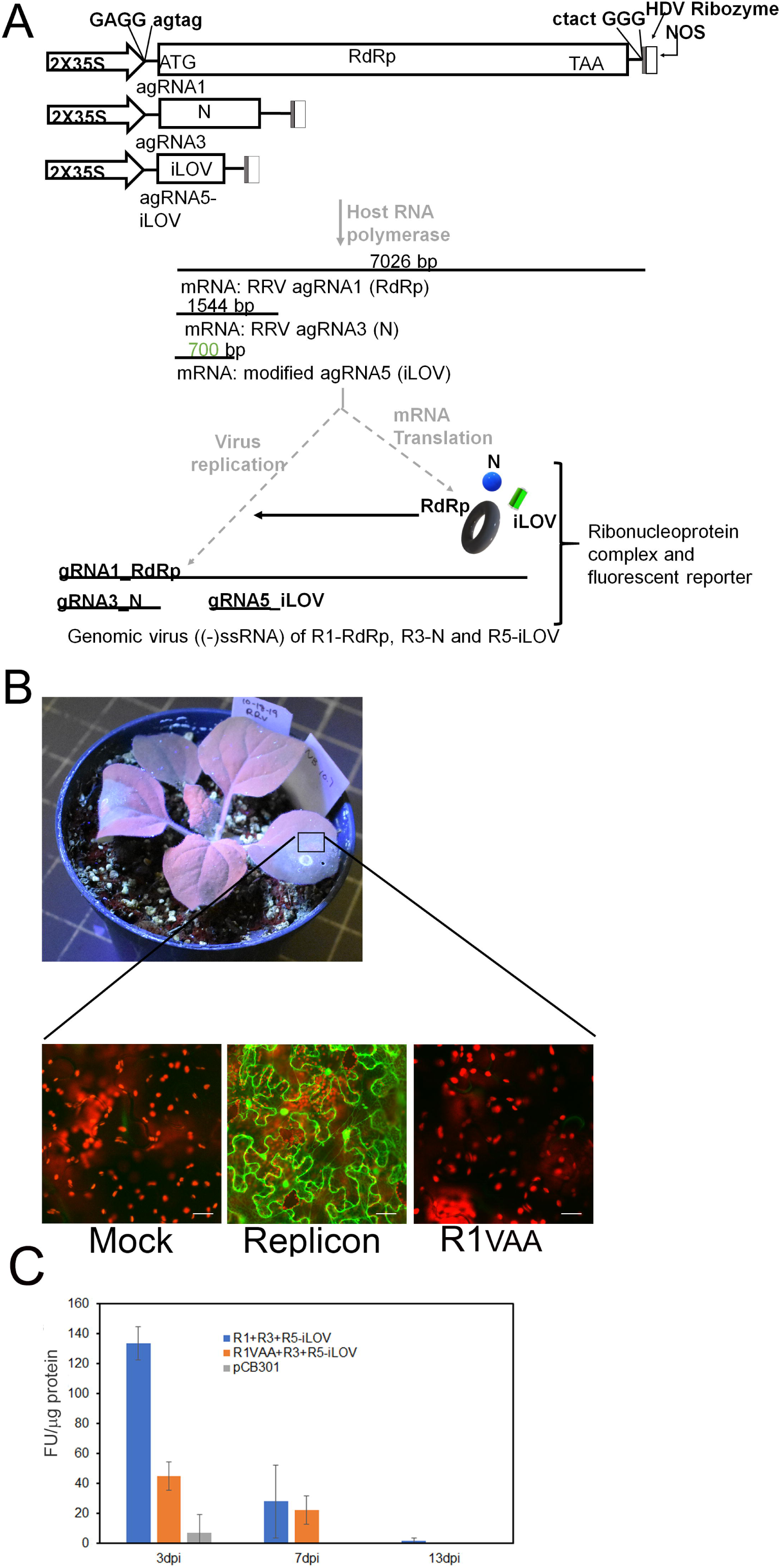
Phylogeny for RNA dependent RNA polymerases of members of the Bunyavirales using representative species provided in the recent taxonomy update reported by Abudurexiti et al., 2019. The tree was generated using IQ-TREE ver. 1.6.11 (76–78) and visualized using iTOL ver 4.4.2 (70). The group containing emaraviruses (including RRV) and related orthobunyaviruses are highlighted in orange. The branches containing tospoviruses and tenuiviruses are highlighted in purple. This phylogeny reveals that the emaraviruses, including RRV, are more closely related to orthobunyavirus species than to tosopviruses. All species used in this phylogeny and Genbank accessions are also provided in Table S1.

For most viruses within the Order *Bunyavirales*, the RdRps contains an N-terminal endonuclease domain and C-terminal polymerase domain (Fig. 2A) (24, 29). The polymerase domain of all negative-strand virus RdRps has six highly conserved motifs known as pre-A, A, B, C, D, and E (29–31). We prepared a multiple sequence alignment using CLC Genomics Workbench 8.0.1 (https://www.qiagenbioinformatics.com/) using the RdRp sequences of 15 viruses belonging to the *Emaravirus* (including RRV), *Orthobunyavirus, Pacuvirus, Lincruvirus, Herbevirus*, and *Shangavirus* genera (Fig. 2B and C). The alignment reveals that RRV shares the linear arrangement of these six active site motifs with members of *Emaravirus* and *Orthobunyavirus* genera. For RRV, the amino acid residues within these motifs are highly conserved with other members of the genus *Emara*- and *Orthobunyavirus*, with only a few similar substitutions in each motif (29–31). The endonuclease domain cleaves RNA, is essential for transcription, and has active site motifs named H, PD, D/E, and T/K (29). These critical active site motifs are represented in the RRV sequences as well as the Emaravirus and Orthobunyavirus members presented in the alignment.

**Fig. 2.**
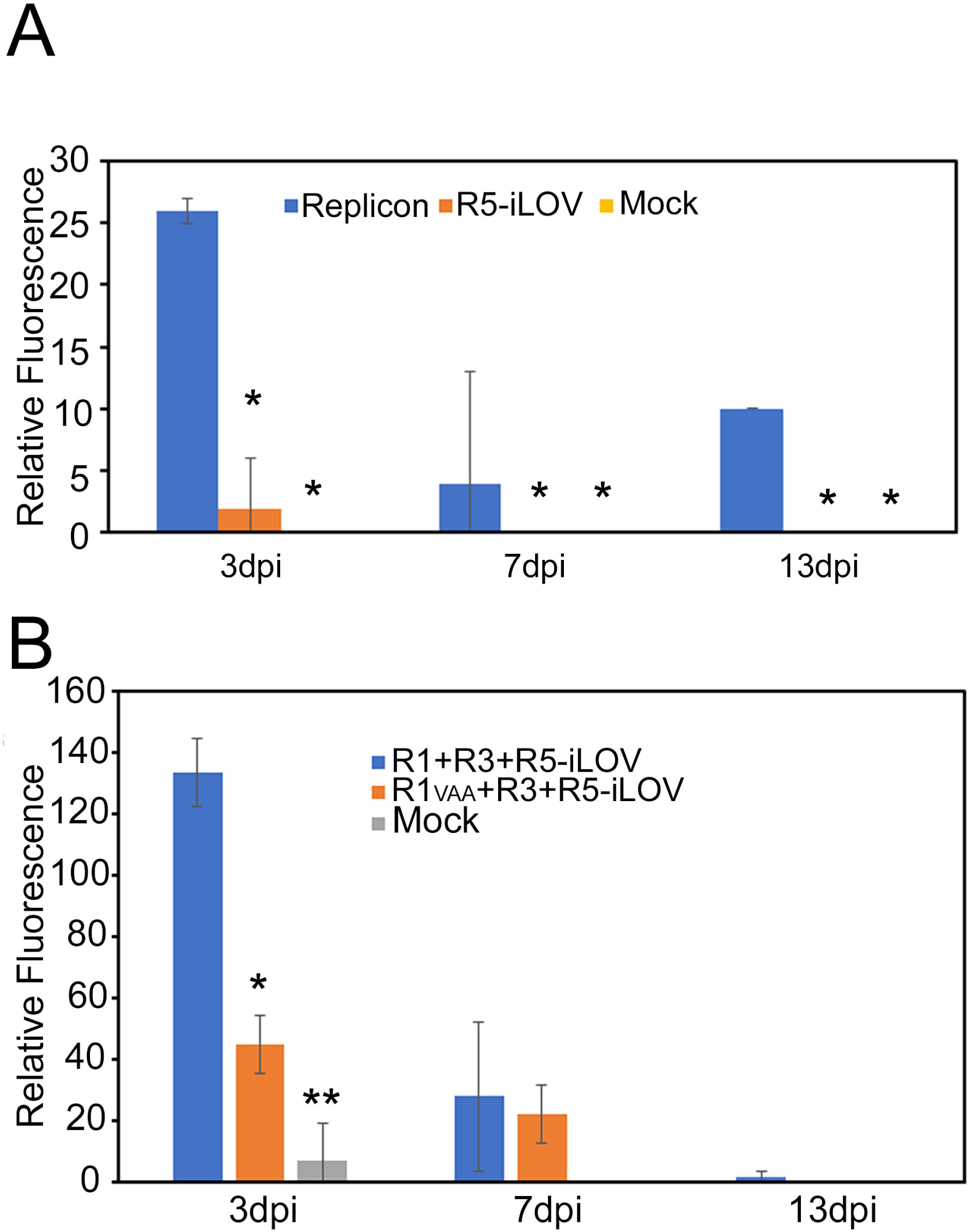
Domain analysis and amino acid sequence alignment of the viral RdRp. Genbank accessions are also provided in Table S1. Protein and domain analysis were carried out using CLC Genomics Workbench 8.0.1 (https://www.qiagenbioinformatics.com/) and by comparing with protein alignments and reported crystallographic structures in the literature (29–31). A. Diagrammatic representation of the RdRp, identifying the endonuclease (EN) domain and the six polymerase active site domains. B. Amino acid sequence alignment of the replicase active sites for fourteen viruses: European mountain ash ringspot-associated virus (EMARaV; YP_003104764), Actinidia chlorotic ringspot-associated virus (ACcrAv; YP_009507925), Redbud yellow ringspot-associated virus (RYRaV; YP_009508083), Pigeonpea sterility mosaic virus 1 (PPSMV; YP_009237282), Rose rosette virus (RRV; YP_004327589), Fig mosaic virus (FMV), Pigeonpea sterility mosaic virus 2 (PPSMV-2; YP_009268863), High plains wheat mosaic virus (HPWMoV; YP_009237277), Raspberry leaf blotch virus (RLBV; YP_009237274), Wēnlǐng crustacean virus 9 (WlCV-9; YP_009329879), Pacui virus (PACV; YP_009666929), Bunyamwera virus (BUNV; AXP32006), La Crosse virus (LACV; AAA62607), Herbert virus (HEBV; YP_009507855), Shuāngào insect virus 1 (SgIV-1; YP_009300681). Green boxes surrounding the letters identifies all identical residues which yellow and gold identify highly conserved residues within each motif. The Bit score identifies the identical and highly conserved residues and is located at the top of each of the aligned motifs. C. Amino acid sequence alignment of the endonuclease using the same species in panel B.

### The N Protein of RRV share similarities with orthobunyaviruses

We constructed a phylogenetic tree of the N protein sequences from 51 species using W-IQ-Tree with 1000 ultrafast bootstraps (28) (Table S2). The predicted RRV N protein clustered with the *Orthobunyavirus* species LACV and BUNV in a common clade (Fig. 3). The closest neighboring cluster were members of the genus *Hantavirus* and *Orthophasmavirus*. The species belonging to the genera *Orthotospovirus* and *Tenuivirus* were distant from the *Emaravirus*, as seen in the RdRp phylogeny.

**Fig. 3.**
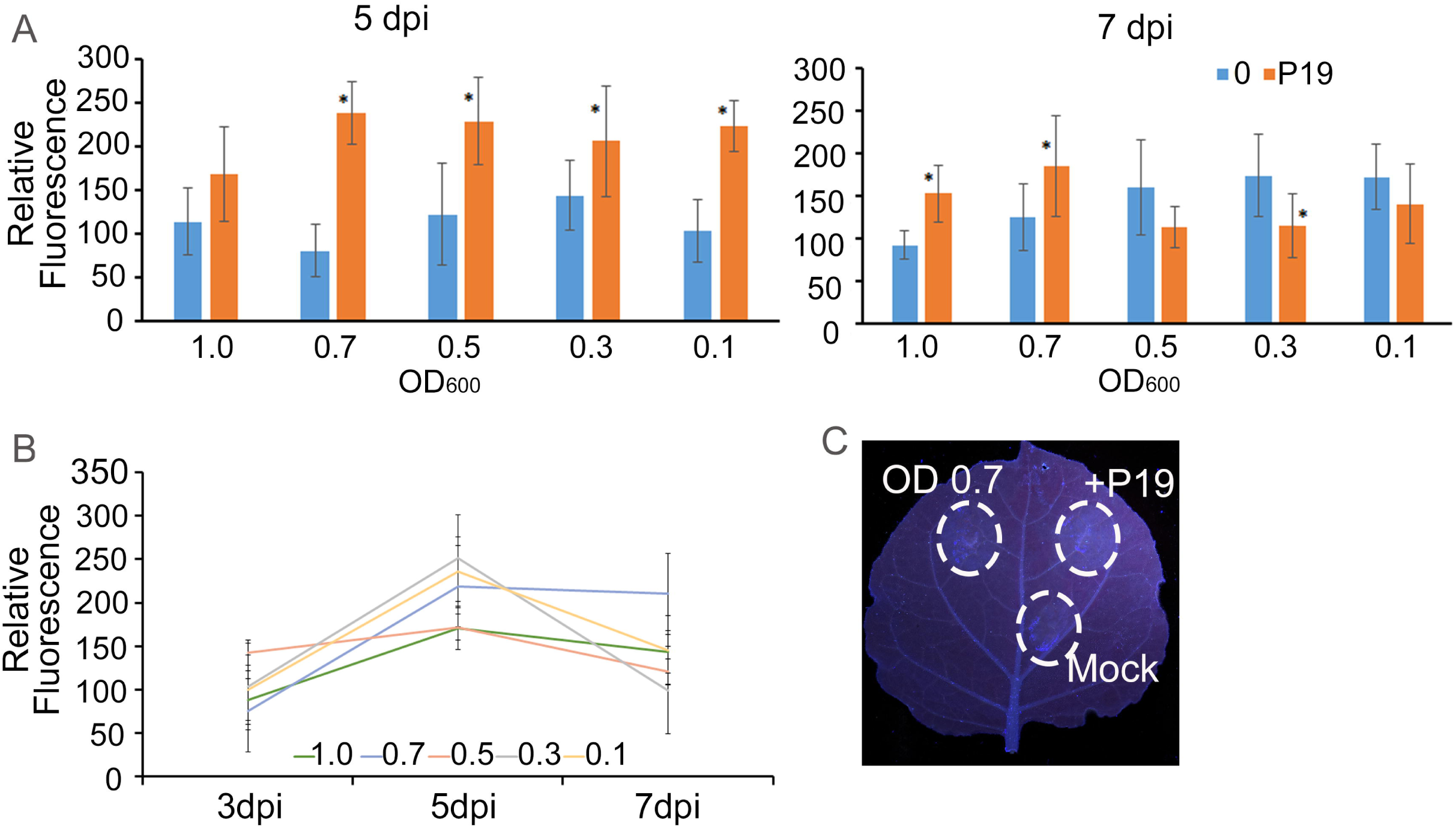
Phylogeny for N protein of members of the Bunyavirales using representative species provided in the recent taxonomy update reported by Abudurexiti et al., 2019. The tree was generated using IQ-TREE ver. 1.6.11 (76–78) and visualized using iTOL ver 4.4.2 (70). Genbank accessions are also provided in Table S2. The group containing emaraviruses (including RRV) and related orthobunyaviruses are highlighted in orange. The branches containing tospoviruses and tenuiviruses are highlighted in purple. This phylogeny reveals that the emaraviruses, including RRV, are more closely related to orthobunyavirus species than to tosopviruses.

The hantavirus, orthobunyavirus, phlebovirus, and tospovirus nucleocapsid proteins were extensively studied using crystallographic, biochemical, and computational methods (24, 32–35). The individual proteins have an N-terminal arm, a C-terminal arm, and a core domain consisting primarily of alpha helices and intervening loops. The RNA binding surface lies in a groove consisting of positively charged amino acids. There are a few β strands in the core domains (32–34, 36, 37). These proteins form higher-ordered oligomers with head-to-tail interactions. To test the hypothesis that the RRV RNA3 encodes the N protein, we used the I-TASSER computational server (38, 39) to predict the 3-D structure and function of the translation product (Fig. 4A and B). The I-TASSER structural prediction was refined using the combined threading and *ab initio* modeling. The top protein models included the nucleocapsids belonging to four orthobunyaviruses: BUNV (PDB:4ijsA), LACV (PDB:4bgpA), *Leanyer virus* (LEAV; PDB:4j1gA), and Schmallenberg virus (PDB:4iduA) (Table 2). The model in Fig. 4A shows an N-terminal coil-coil arm and a C-terminal helical arm that extends from the core, similar to many orthobunyavirus N structures. The overall predicted RRV N protein has eight predicted helical strands. The C-score for the model is −3.91, which is in the value range of −5 to 2 for a confidence estimate. The TM score is a measure of topological similarity between the protein and template models, which is 0.29 ±0.09. The statistical values are near the lower end of the acceptable ranges to validate the models due to the low amino acid sequence identity with the templates (24).

**Table 2.** Total proportion of systemically infected plants confirmed by RT PCR using agRRV4 F1/R1 primer set

| Constructs | *N. benthamiana* |
| --- | --- |
| Rose sap inoculum | 12/12 |
| Buffer treated | 0/8 |
| pCB301 | 0/12 |
| RRV1-4 | 16/18 |
| RRV1-7 | 17/18 |
| 7R5-iLOV | 8/8 |
| 4R2-iLOV | 8/8 |
| 7R2-iLOV | 8/8 |

The proportion of infected *N. benthamiana* were pooled from two experiments. All plants were assayed by RT-PCR at 25 dpi.

**Fig. 4.**
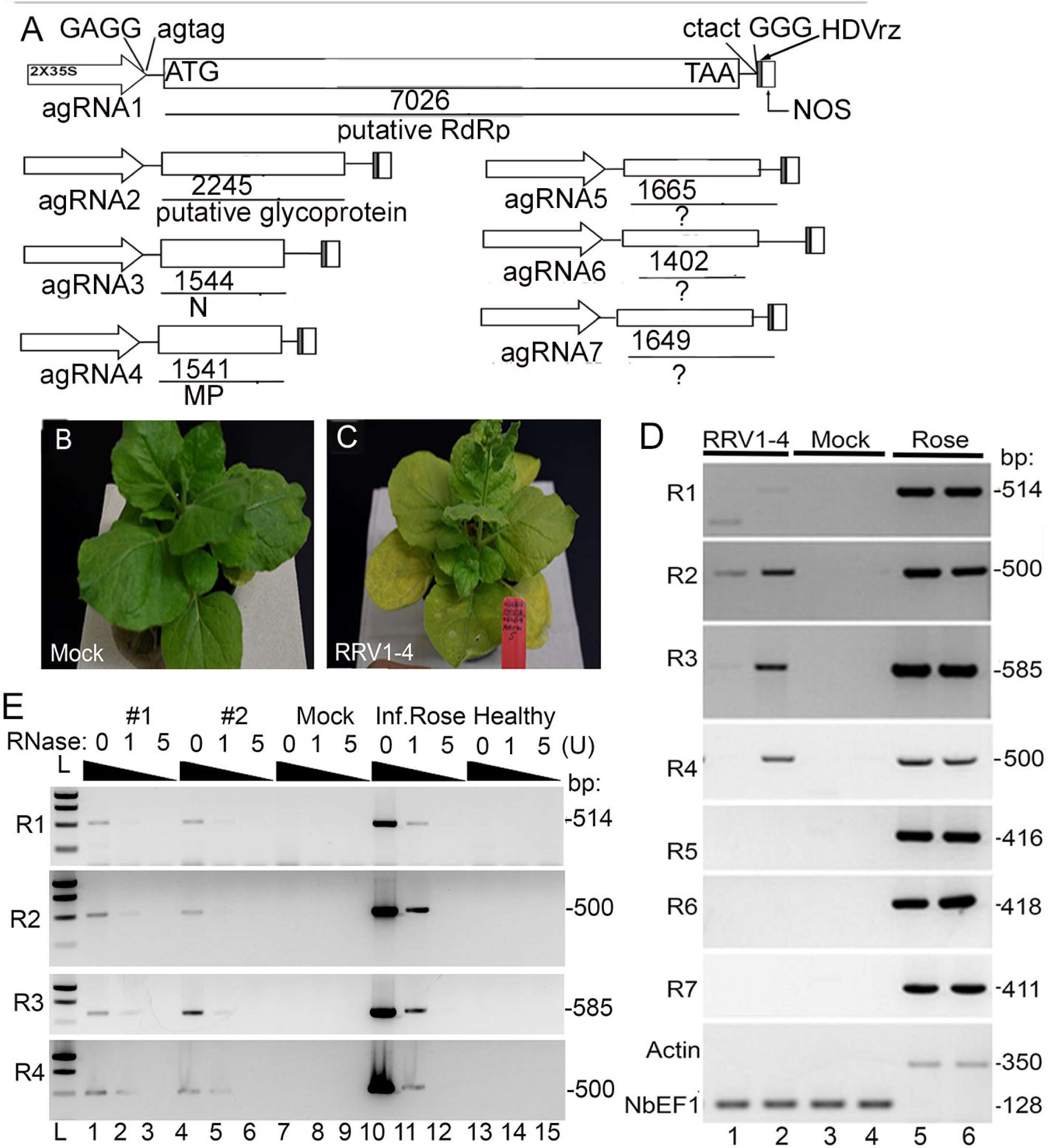
Three dimensional models and sequence alignment for the N protein. The models were generated and analyzed using I-Tasser and The PyMOL Molecular Graphics System 2.0 (Schrodinger, LLC). Proteins were aligned for domain analysis using CLC Workbench 8.0.1. A. The 3D model of the RRV glycoprotein. The folding domains are colored, and the locations of the N- and C-termini are identified. B. Model showing RNA binding cleft of the N protein. The orange ribbon represents RNA within the binding cleft. C. Amino acid sequence alignment of emaravirus N proteins generated using CLC Biology Workbench. The region represented here encompasses the RNA binding region and highlights the consensus with 100% identity the alignment. There is a stretch of highly conserved residues between positions 140 and 280.

Ligand binding sites were predicted using COFACTOR and COACH and built on the I-TASSER structure and graphically presented using PyMOL to demonstrate the core alpha-helical region contains the RNA-binding cleft (Fig. 4B and C). The closest models for this regional interaction are the BUNV N protein (PDB: 3zlaA) and the LEAV N protein (PDB: 4ji1ja). The C-score range for ligand binding was 0.29 and 0.08, which is within the score range of 0-1 (Table 2). Since positively charged amino acid sidechains mediate hydrogen bonds with the RNA phosphate backbone, we identified highly conserved Arg, Asp, Glu, Lys residues between amino acid positions 100 to 312 that are likely involved in RNA interactions.

### The terminal untranslated regions folding of RRV genomic RNA is similar to orthobunyaviruses

We mapped the UTRs for each RNA segment, and genomic 5’ UTRs are generally longer than the 3’ UTRs except for RNA1, which has a 3’ UTR of 107 nt and a 5’ UTR of 88 nt. For RNA-2 through −7, the 5’UTRs ranged from 190 to 631 nt in length while the 3’ UTRs were between 50-99 nt in length (Table 1). Notably, the RNA1 has shorter UTRs than the other segments.

**Table 1.** Genome characteristics of Rose rosette viruses

| RRV Genomic segments | 3' UTR (nt) | Protein amino acid length | 5' UTR (nt) | Potential Function<sup>a</sup> |
| --- | --- | --- | --- | --- |
| segment 1 (7026 nt) | 107 | 2276 | 88 | RNA-dependent RNA-polymerase |
| segment 2 (2245nt) | 50 | 645 | 257 | Glycoprotein |
| segment 3 (1544 nt) | 99 | 319 | 494 | Nucleocapsid |
| segment 4 (1541nt) | 83 | 361 | 372 | Movement protein |
| segment 5 (1665 nt) | 61 | 467 | 200 | unknown function |
| segment 6 (1402 nt) | 68 | 233 | 631 | unknown function |
| segment 7 (1649 nt) | 61 | 465 | 190 | unknown function<sup>b</sup> |

<sup>a</sup>Protein and domain analysis carried out using InterPro and HMMER (EMBL-EBI) and BLASTP (GENBank)
<sup>b</sup> The high plains wheat mosaic virus (HPWoMV) is also a member of the genus *Emaravirus* and encodes eight genome segments. RNA7 and RNA8 encode silencing suppressor proteins (Gupta et al. 2019). RRV encodes seven (not eight) segments. The HPWoMV and RRV P7 proteins share no real sequence conservation to infer similar functions.

We compared the terminal untranslated regions (UTRs) of each RRV genomic segment with the UTRs of BUNV (Fig. 5)(29) based on prior reports of essential base pairing between the 3’ and 5’ UTRs BUNV terminal nucleotides for successful replication and transcription (38, 39). For RRV, the predicted 5’ UTRs were generally longer than the 3’ UTRs, except for RNA1, which had 88 bp at the 5’ end and 107 bp at the 3’ end. Using the mFOLD Web Server (40), we obtained the RNA folding results which revealed that the first twenty nucleotides of the 3’ and 5’ terminal regions are complementary, with only seven nucleotide differences among the genome segments (Fig. 5). Remarkably, the terminal seven nucleotides of the RRV genome segments are well conserved with the terminal seven nucleotides of the BUNV segments S and L (41, 42). According to Barr et al., (2005), the 5’ UTR sequence on the BUNV S segment includes a stretch of five bases, ACUAC, that is essential for transcription. This element is not present in the UTRs of any RRV segments. The combined computational analyses indicate that RRV RdRp and N proteins, along with the UTRs, share essential similarities with orthobunyaviruses making these useful models for predicting the general features of RRV transcription and RNA replication. Therefore, we considered the integrity of these highly conserved features when developing a new system to study RRV mRNA synthesis and encapsidation at the cellular level, as well as virus-host interactions at the whole plant level.

**Fig. 5.**
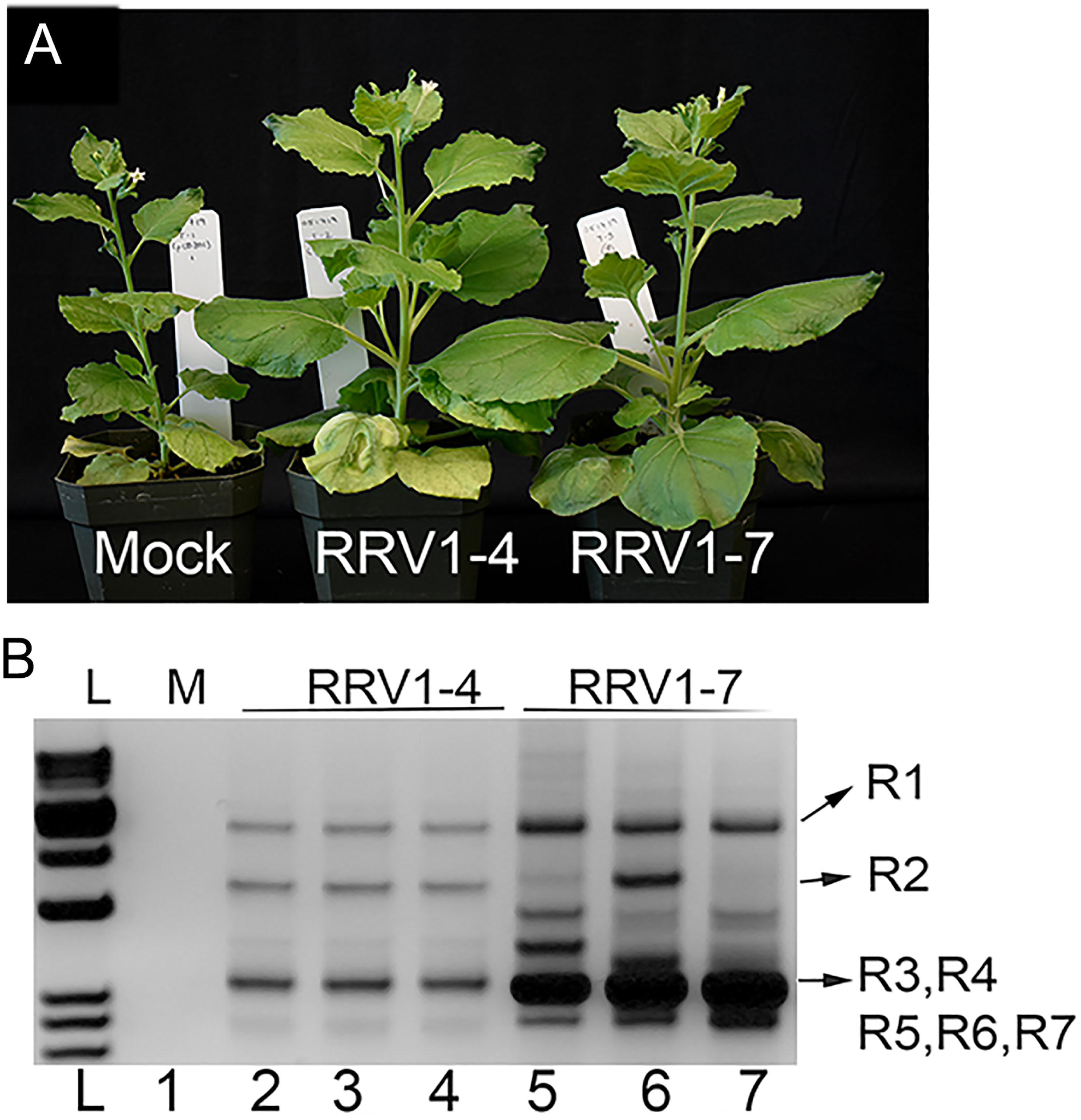
Results of mFOLD showing hybridization of the 3’ and 5’ ends of each RRV and BUNY segment. The initial ΔG for RRV RNA1 through RNA7 is −43.2, −161.30, −138.0, −341.90, - 137.50, −125.00, and −129.40 respectively. The ΔG for BUNV L, M and S segments are - 131.50, −106.50, and −128.60 respectively. There are two segments that share a remarkable identity. First is the terminal seven (AGUAGUG) bases of all seven RRV segments and with the BUNV segments shown here. A significant difference is the mismatch of UU in rose rosette is at the same location as a single mismatch A at the 3’ end of BUNV RNAs. This is followed by CUCC, which is identical among all RRV segments and BUNV RNA1 and RNA3. For RRV, the seven segments either share a common CUCCCU/A, CUCCUC/U.

### Successful insertion of the fluorescent protein iLOV into RNA5 as a reporter for virus replication in *N. benthamiana* leaves

We developed a reporter mini-replicon system that can be delivered by agroinfiltration to *Nicotiana benthamiana* leaves for *in vivo* studies. Two cDNAs encoding the antigenomic RNA1 (7026 bp) and RNA3 (1544 bp) encoding the predicted RdRp and N proteins, respectively, were introduced into binary plasmids between a duplicated 35S promoter and a nos terminator. The terminal 5’ A of the agRRV cDNA was fused to the 3’ G residue of the promoter (Fig. 6A) (18). The agRRV cDNA 3’ end was fused to the HDRz. To examine the template activities of these combined viral proteins, we prepared a third plasmid derived from the cDNA of agRNA5 (1665 bp). We replaced the short viral open reading frame with the photoreversible fluorescent iLOV protein as a reporter for virus replication. Thus, the RNA5-iLOV binary construct contains the iLOV coding sequence flanked by the RRV 3’ and 5’ UTRs. The iLOV protein is a 10 kDa flavin-based alternative to GFP, derived from plant phytotropin, and was reported to be an excellent alternative to GFP for monitoring virus infection (17). Agrobacterium cultures harboring these plasmids were combined in equal ratios and introduced by infiltration into *Nicotiana benthamiana* leaves. At 2-days post infiltration (dpi) leaves were examined using a hand-held UV lamp, and there was no apparent fluorescence. However, using a confocal microscope at 3 dpi, we observed iLOV fluorescence in neighboring epidermal cells (Fig. 6B). The fluorescence was primarily in the cytoplasm and nucleus as expected. The mock-treated leaves showed no fluorescence.

**Fig. 6.**
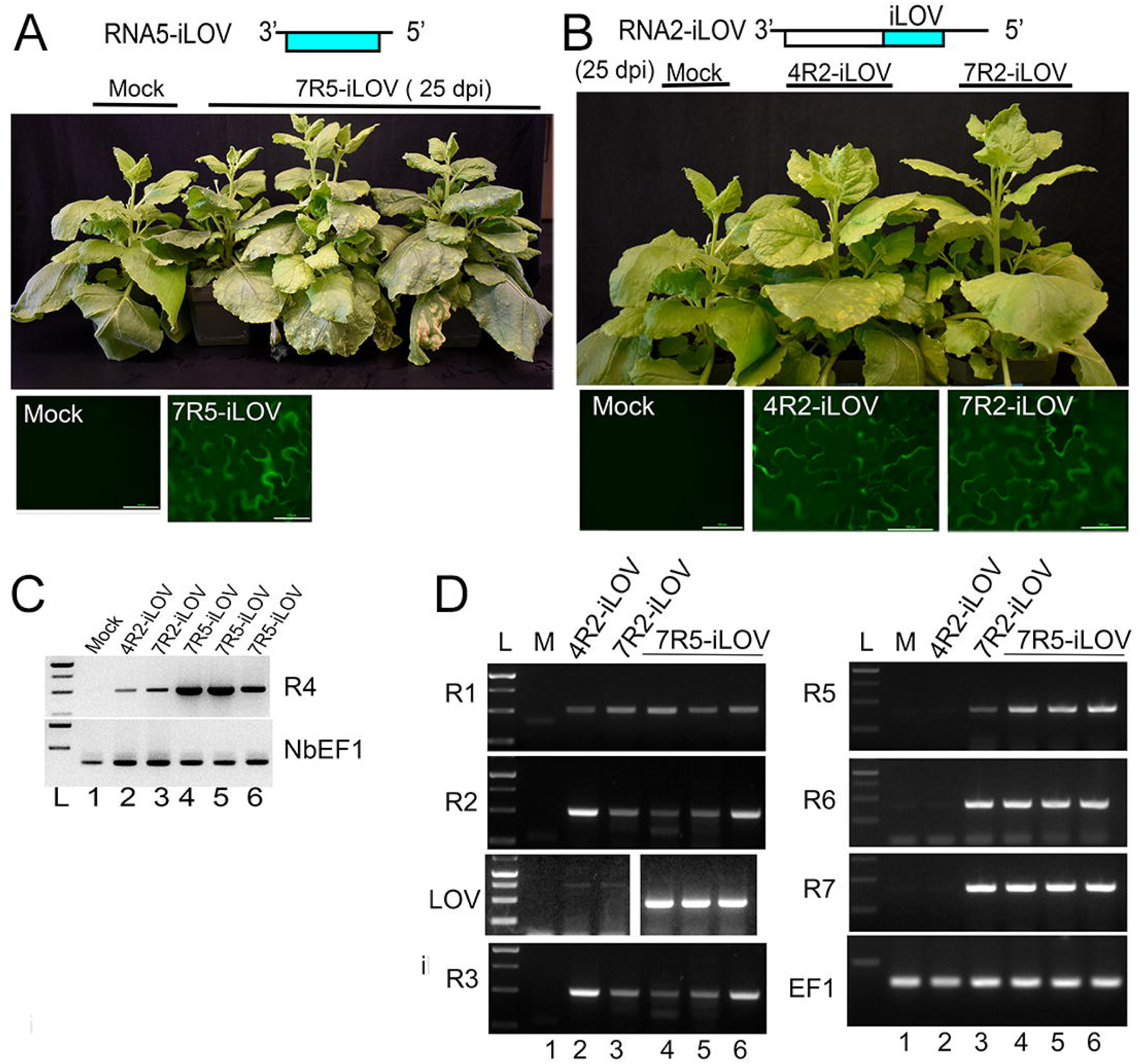
RRV minireplicon system using R5-iLOV as a template. A. Diagrammatic representation of the RRV anti-genome replicon. The duplicated 35S promoter is identified by an arrow. The GAGGAGTAG sequence is identified to show the pent-ultimate residue of the promoter is flush with the first residue of the RRV genome. The HDRz is at the 3’ end of the antigenome sequence and functions upon transcription to cleave the end to produce an exact terminal residue for the virus agRNA segment. The predicted names for each coding region are identified in the open boxes: RNA dependent RNA polymerase (RdRp), nucleocapsid (N), and iLOV. After the agrobacterium delivery of binary vectors to the nucleus, there is a single round of transcription led by the host RNA polymerase to produce an antigenomic mRNA that produces the RdRp, N and iLOV proteins. These can act on the same antigenomic mRNA to initiate reverse transcription to produce genomic RNA (gRNA) from each of the segments as the start of replication. Continued replication results in increasing levels of iLOV expression. B. Infiltrated leaves were harvest for confocal imaging at 2 and 3 dpi. The iLOV fluorescence is seen in the cells containing the functioning minireplicon system, but not in the mock-treated cells or leaves treated with R1(VAA) + R3+R5-iLOV. Scale bars represent 20 µm. C. The average Fluorescence Units (FU) per ug of protein (n=3) was reported at 3 dpi following agrodelivery of the RRV R1, R3, and R5-iLOV; the R1VAA, R3 and R5-iLOV; or pCB301 backbone without iLOV. Raw data was normalized to the fluorescence level of a healthy leaf sample, and mock treatments using Microsoft Excel.

The iLOV expression could have been the product of primary transcripts produced from the plasmids or the result of RNA synthesis by the viral encoded RdRp. To determine if the viral RdRp was responsible for iLOV expression, we engineered a mutation into the RNA1 segment to disrupt the viral polymerase activities. The C motif of the RdRp contains the catalytic domain of the polymerase. Substitution mutations of the conserved Ser-Asp-Asp tripeptide are predicted to hamper RNA polymerase activity or alter the polymerase metal cofactor preferences (32, 38). In this study, we replaced the Ser-Asp-Asp tripeptide of the RdRp with a Val-Ala-Ala tripeptide to produce RRV RNA1_VAA_. Using confocal microscopy at 2 or 3-dpi, we did not see iLOV fluorescence in leaves expressing RNA1_VAA_, RNA3, and RNA5-iLOV (Fig. 6B).

Fluorometric analysis was conducted using the supernatants of crude homogenates (derived from infiltrated leaves) at 3 dpi. The average fluorescence units (FU) per µg of protein (n=3) for each treatment was normalized with the healthy leaf homogenates (Fig. 6C). Fluorescence due to the RNA1, RNA3, and RNA5-iLOV was 4-fold higher than fluorescence detected in supernatants from samples expressing RNA1_VAA_, RNA3, and RNA5-iLOV, or mock samples that were agroinfiltrated to deliver the empty pCB301 plasmid (Fig. 6C). These combined data indicate that RNA1 and RNA3 are the minimum segments required for iLOV expression from RNA5 and that RNA1 likely encodes the viral RdRp.

### Characterization of the RRV surface glycoprotein

The bunyavirus RNA2 (or M segment for some families) encodes a polyprotein precursor that is cleaved into two surface glycoproteins known as Gn and Gc. The Gc identifies cell surface receptors for virus internalization and subsequent virion disassembly in the cytoplasm (43–45). During virus replication and protein synthesis, the polyprotein precursor inserts into the ER through the signal recognition particle. The signal peptide is cleaved before maturation cleavage to produce the Gn and Gc proteins. Most bunyavirus glycoproteins contain Golgi localization signals and the enveloped virions bud through the Golgi apparatus (46, 47). The RRV glycoprotein precursor was aligned with other emaravirus polyproteins (Fig. S1) to identify conserved elements and essential features presented in Fig. 7A. Three-D modeling was conducted using I-Tasser and Pro-Origami (38, 48) and these outputs were compared with the conserved sequence motifs identified in the literature for bunyavirus glycoproteins (45, 49). The N-terminal 19 amino acids contain the signal peptide (S) for ER insertion. There are three transmembrane domains located between amino acid positions 107-137, 178-193, and 587-609. In the Gn region, between the two transmembrane domains are two adjacent Golgi retention signals, R(K)xD and F(Y)Y. A third potential Golgi retention signal lies near the C-terminus of the polyprotein. There is a potential ER retrieval motif (KKXX) at amino acid position 287 and potential ER export motif Y(F)Y (47). RRV and several other emaraviruses have a putative proteolytic cleavage site, Ala-Lys-Ala-Asp-Asp adjoining the second transmembrane domain and the putative ER export site DxE(Q) (50, 51). The more extended Gc region has two potential ER export sites (KK and Y(F)Y) and a potential caspase 3 cleavage site (FVDSS). The RRV glycoprotein has a conserved GCYDCQxG motif that resembles the GCYDCQNG motif that present in the phlebovirus glycoprotein (Fig. 7A)(50, 52, 53).

**Fig 7.**
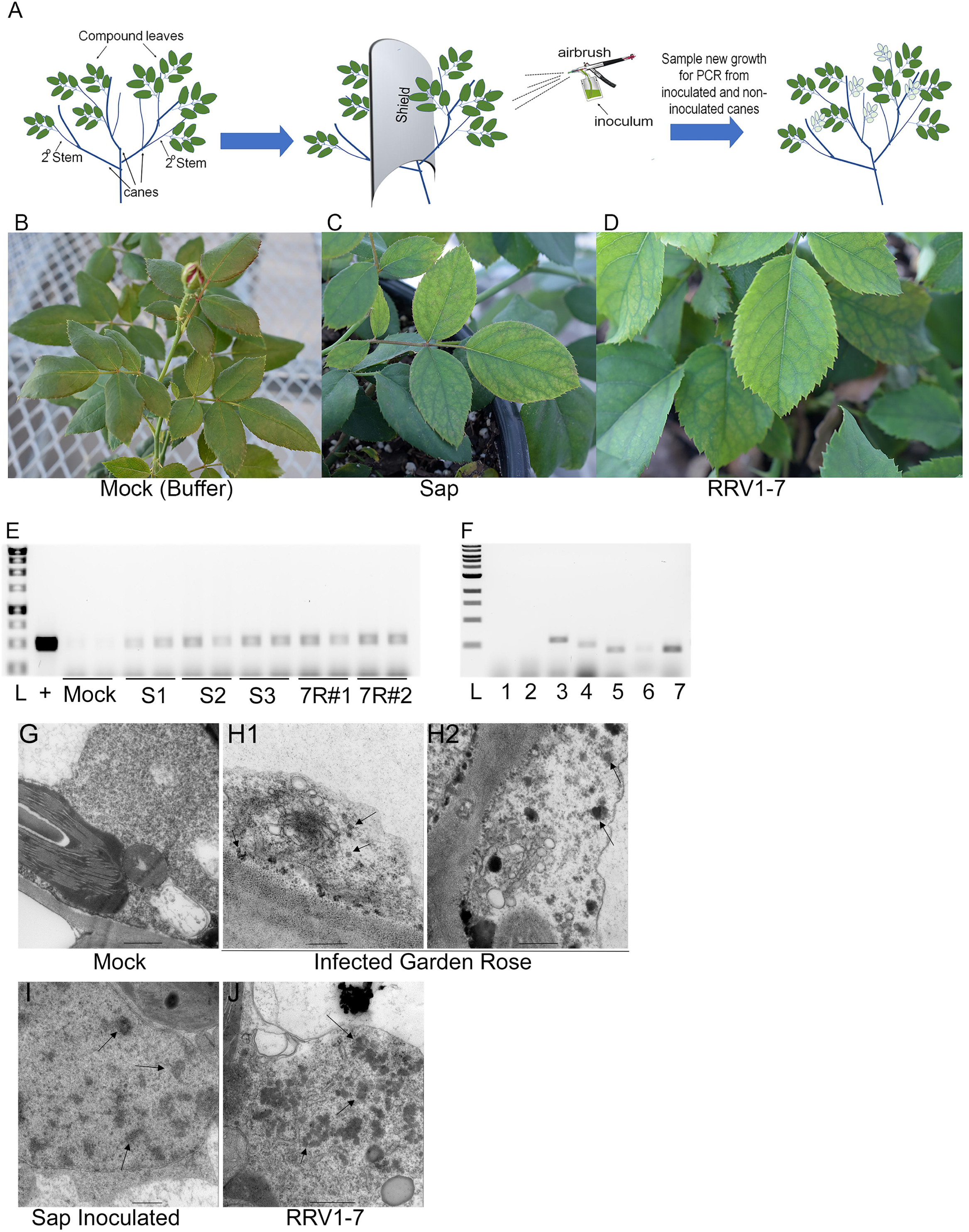
Alignment and model predictions of the RRV glycoprotein. A. Diagrammatic representation of the R2 encoded polyprotein with key functional domains identified. The signal recognition particle (SP) is followed by two transmembrane domains (TM) and two potential ER export signals, a potential Caspase 3 cleavage site followed by another TM, and a C-terminal Golgi retention signal. The positions of each of these domains are identified below the bar. The sequence alignment surrounding the second TM is featured below. This region contains two potential Golgi retention signals and an ER export site adjacent to a predicted cleavage site that is conserved among emaraviruses and other orthobunyaviruses. Proteolysis at this location produces a smaller membrane-associated glycoprotein that is retained in the Golgi. The C-terminal longer glycoprotein also has ER export sites and Golgi retention signals, suggesting that these two glycoproteins co-localize in the Golgi. B, C. I-Tasser generated 3D model of the RRV glycoprotein precursor. Domains are colored and the locations of the N- and C-termini are identified, as well as the three TM domains and the Gc motif. The same model is presented in panel B and C but from different angles. D. Pro-Origami depiction of the 3D model. The parallel and antiparallel B-sheets are surrounded by blue boxes. The 20 conserved Cys residues are identified in the structure. These stabilize the folded protein.

For orthobunyaviruses and hantaviruses, the Gc is a class II fusion protein. The X-ray and crystal structures of the Gc from *Rift valley fever virus* (RVFV), *Hantaan virus* (HNTV), *Puumala virus* (PUUV) provide significant insight into the pre- and post-fusion structures that exist between the Gc trimers of vertebrate and invertebrate viruses (43, 54, 55). We examined the I-Tasser generated model of the RRV glycoprotein precursor for features of a fusogenic glycoprotein. The predicted 3D structure of the RRV glycoprotein precursor had a C-score of −1.75 and an estimated 12.1Å resolution (Fig. 7B and C). The top ten threading templates used by I-Tasser included the post-fusion conformation of the GC proteins of *Thrombocytopenia syndrome virus* (a phlebovirus; PDB: 5G47), the RVFV (an orthobunyavirus; PDB: 4HJ1), and the NMDA receptor ion channel (PBD: 4pe5B)(Table 2). The detailed predicted 3D folded structure of the glycoprotein precursors resembled the I-Tasser predicted structure reported for emaravirus *Pigeonpea sterility mosaic virus* (PPSMV-1) but was dissimilar from the class II fusion proteins of HNTV, and PUUV (44, 51). These orthobunyavirus class II fusion proteins have three domains of antiparallel β-sheets. The N- and C-terminal β-strands are in proximity.

The topological diagram (Fig. 7D) identifies two regions of primarily short parallel β-strands and three domains with antiparallel β -strands connected by stretches of α-helices and other helices. There is a GCYDCQxG sequence that resembles the GC-x2-C sub motif that is found on an exposed loop of the phlebovirus glycoprotein and is characteristic of class II fusion proteins (Fig 7C and 7D).

The folding domains and the trimeric conformations of other bunyavirus glycoproteins are supported by disulfide bonds based on 10 or more conserved cysteine (Cys) residues (43). We examined the pattern of residues within the predicted 3D model to identify disulfide bonds that could stabilize the folded protein and motifs that govern membrane fusion. We aligned twelve emaravirus glycoprotein precursors (Fig. S1) and identified 20 conserved Cys residues, which we then located in Fig. 7D on the topological diagram. RRV has two additional Cys residues that are not conserved with other bunyaviruses (Fig. S1). Also, there is a GCYDCQxG sequence that resembles the GC-x2-C sub motif that is found on an exposed loop of the phlebovirus glycoprotein and is characteristic for class II fusion proteins (Fig 7C and 7D).

Most bunyavirus glycoproteins have a conformational “cd loop,” also known as the “target membrane-interacting region.” This region contains a WG-X2-C, WG-X3-C, WG-X5-C, or WR-x5-C motif. We identified the W_102_ and W_119_ in RRV are highly conserved among all emaraviruses. A W_110_ is conserved among five emaravirus species but does not reside within the WG(R)-Xn-C motif that is predicted for insertion into target membranes for membrane fusion (Fig. S1). For some viruses, the class II fusion machinery uses His residues as a pH sensing mechanism that recognizes changes in the environmental pH of the endosome. The His162, His205, and His218 are highly conserved among emaraviruses, although their role in pH sensing needs further assessment. These combined data highlight significant similarities between the glycoprotein of emaraviruses and bunyaviruses.

### RRV infectious clone produces systemic infection in *N. benthamiana* plants

We prepared full-length infectious clones of all RRV segments. The binary plasmids contain the full-length cDNAs for agRNA1 (7026 bp), agRNA2 (2245 bp), agRNA3 (1544 bp), agRNA4 (1541 bp), RNA5 (1665 bp), agRNA6 (1402 bp), and agRNA7 (1649 bp)(18)(Fig. 8A). All antigenomic cDNAs were positioned immediately adjacent to the upstream CaMV 35S promoter and downstream HDRz to produce viral transcripts with authentic 5’ and 3’ ends (Fig. 8A). All constructs were confirmed by sequencing before transformation into *A. tumefaciens* (Fig. 8A).

**FIG. 8.** *N. benthamiana* infected with the infection clones RRV1-4 and RRV1-7. A. Diagrammatic representation of the infectious clone. As in Fig. 6, open arrows indicate the duplicated 35S promoter. Adjacent sequence directly fusing the 5’ end of the antigenomic cDNA to the promoter is highlighted above the arrow. Each open box represents the open reading frame and is identified as the putative RdRp, glycoprotein, N, movement protein (MP), and unknown (?) proteins. The C terminal sequence fused to the HDRz and Nos terminator is indicated. The iLOV sequence is highlighted in blue. Below the R2-iLOV is a diagram demonstrating the polyprotein is cleaved into two products. The molecular weights are provided. B, C, D. Images of mock-treated, RRV1-4, and RRV1-7 infected plants at 25 dpi. E. RT-PCR verified accumulation of RNA1 through RNA4 in two samples of *N. benthamiana* (lanes 1, 2) and control rose samples (lanes 5, 6). Two mock control samples were included (lanes 3, 4). RNA segments are identified on the left. The amplicon sizes are identified on the right of each ethidium bromide-stained 1.2% agarose gel. F. Multiple RT-PCR using three RNA samples from plants infected with RRV1-4 (lanes 2-4) and three samples from RRV1-7 (lanes 5-7). One RNA sample from mock-treated plants (M, lane 1) was included. Amplicons representing each RNA1 through 7 are identified on the right. Narrow arrows point to individual products while the block arrow identifies the aggregate of bands representing RNAs3-7, which co-migrate on a gel. The “*” identifies intermediate size amplicons, as reported in Babu et al. (2016)(79). G. RNAse protection assay detecting double-stranded RNAs. Total RNAs were treated with DNAse I and then treated with 0, 1, and 5 units (U) of RNase I to digest single-strand RNA, leaving dsRNA intact. RT-PCR amplification is carried out using the same primers as in panel D and listed in Table 1. Products were analyzed by 1.2% agarose gel electrophoresis. The RNAs 1 through 4 are identified on the left, and the size of each amplicon is indicated on the right. L= 1kb ladder.

**FIG. 8.** *N. benthamiana* infected with 4R2-iLOV, 7R2-iLOV and 7R5-iLOV. (A) iLOV fused to ORF2 on RNA2 and introduced into RNA5 replacing the ORF5. Images of plants at 25 dpi that are mock-inoculated or systemically infected with RRV 4R2-iLOV and 7R2-iLOV. Epifluorescence microscope images of infected leaves at 13 dpi shows fluorescence in epidermal cells. Bars= 100 µm. (B) Images of plants at 25 and 35 dpi that are mock-inoculated or systemically infected with RRV 7R5-iLOV. Epifluorescence microscope images of infected leaves at 13 dpi shows fluorescence in epidermal cells. Bars= 100 µm. (C) RT-PCR diagnostic using agRRV4-F1/R1 primer set to identify infected samples. PCR primers are detecting EF1 as internal control showing equal loading in each lane. L=1 kb ladder. (D) RT-PCR detecting each viral segment identified on the left. Samples infected with 4R2-iLOV, 7R2-iLOV and 7R5-iLOV are identified above the gels. EF-1 is a control PCR showing equal loading in each lane.

Typically, systemic plant virus infection requires the activities of the viral replicase, virus movement protein, and encapsidation factors. While we have bioinformatically and experimentally identified the RdRp, N, and glycoprotein, a previous report described the RRV RNA4 to encode the virus movement protein (56). We examined whether RNA1 through RNA4 were sufficient to produce systemic infection or if all seven RNA segments are required for systemic infection. We agro-delivered plasmids encoding RRV RNA1 through RNA4 (RRV1-4) and RRV RNA1 through RNA7 (RRV1-7) to *N benthamiana* plants. *Agrobacterium* cultures containing each virus cDNA were combined in equal ratios and then infiltrated to young *N. benthamiana* leaves. These infectious clones did not contain the iLOV reporter. Plants were mock-inoculated using *Agrobacterium* containing the backbone binary vector. *N. benthamiana* plants were also rub inoculated using sap from an infected rose plant as a positive control for infection (Table 3). These plants were monitored for visual symptoms for up to 25 dpi. The symptoms ranged from mild to significant leaf yellowing (Fig. 8B, C, and D).

**Table 3.**
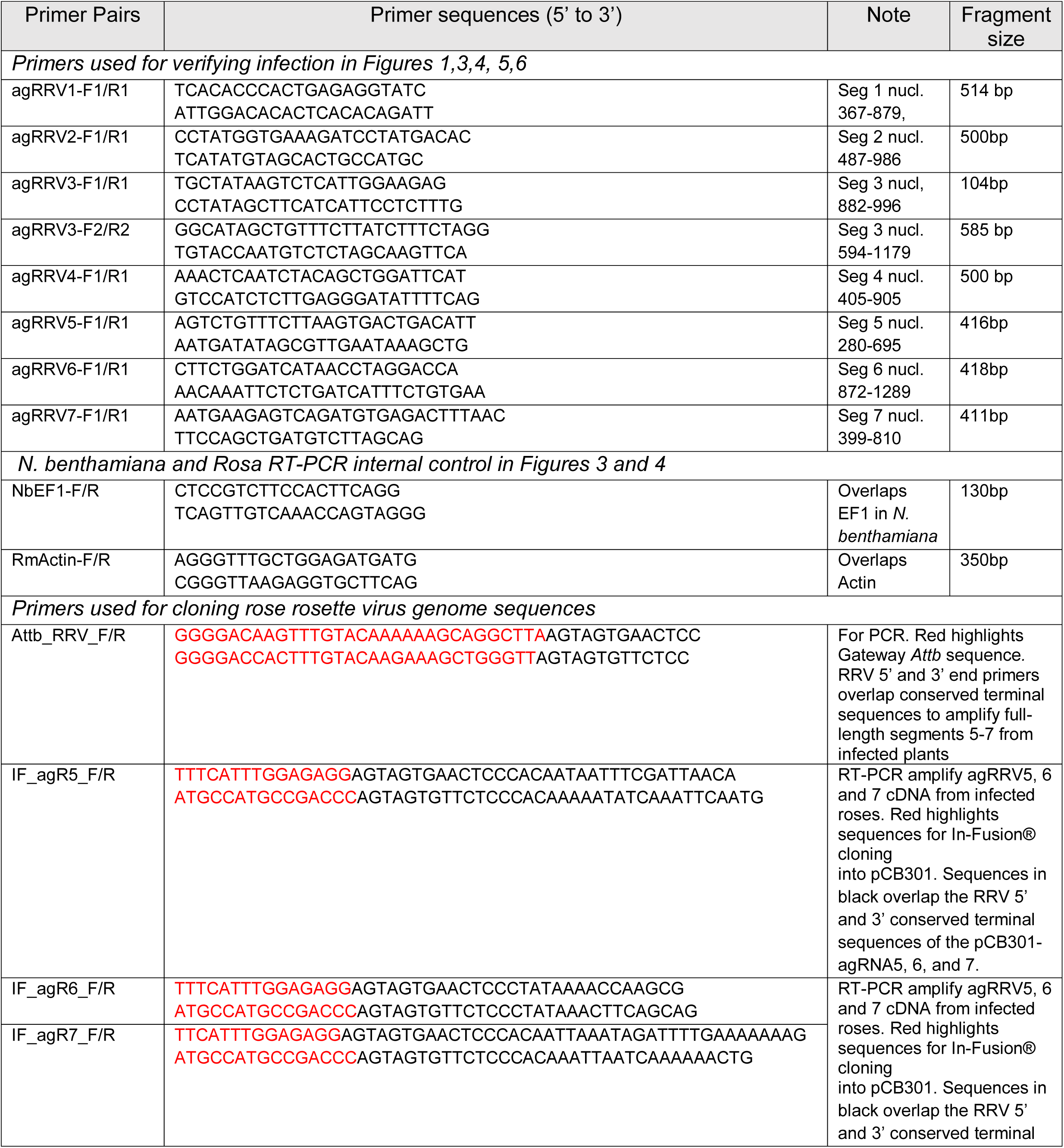
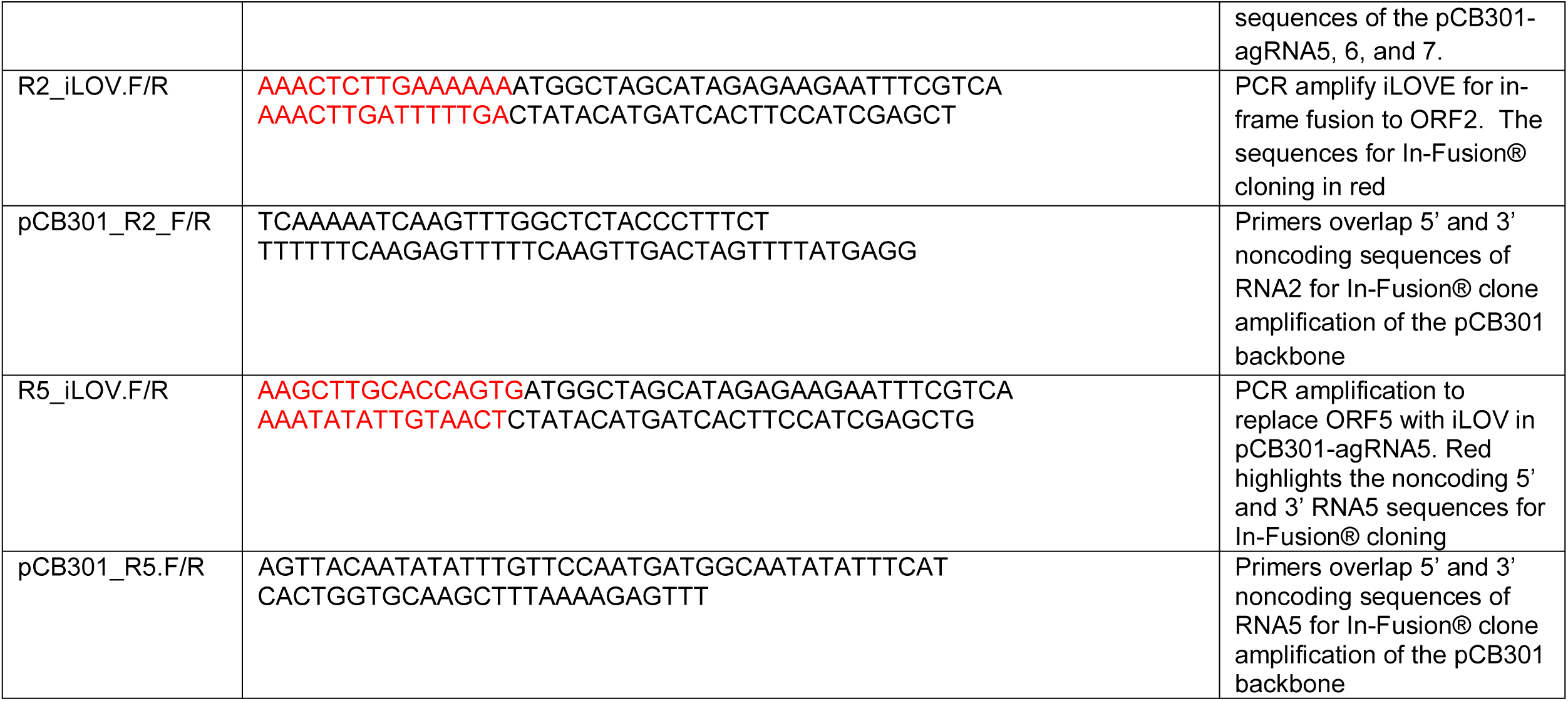
Primers for RT-PCR verification of virus infected plants and cloning

At 25 dpi, we harvested RNA from upper non-inoculated leaves and conducted RT-PCR using primers to detect RNA4 transcripts and antigenomic RNA. All sap-inoculated plants were systemically infected (12/12), 75% of plants treated with RRV1-4 and 85% of plants treated with RRV1-7 were infected. These data suggest that RRV1-4 contains the replication proteins and MP needed to infect plants systemically. For plants infected with RRV1-4, we recovered 5 µg of total RNA from each plant and conducted individual RT-PCR to verify the presence of each RNA segment (Fig. 8E). As an internal control, we used primers to detect the translation elongation factor 1 (EF1) in *N. benthamiana*. Equal volumes of PCR products were loaded in each lane. PCR products detecting RNA1 through RNA4 sequences were amplified, and no other contaminating viral RNAs were detected. These PCR products were sequenced to verify they originated from RRV. Notably, the amplicon levels representing RNA1 through RNA4 varied between samples. PCR products of RNA1 routinely accumulated much less than the PCR products of RNA2, RNA3, and RNA4 using these primer sets. As an external control, similar RT-PCR was carried out using RNA from RRV infected roses. Primers are detecting *Actin* as an internal control. We recovered PCR products representing RNA1 through RNA7 from these infected plants.

In another set of experiments, we conducted multiplex RT-PCR to detect all segments in RRV1-4 and RRV1-7 infected plants simultaneously. These assays employed the aatbRRV F/R primer set (Table 4) that recognizes the conserved 5’ and 3’ sequences and produces amplicons of all seven RNAs in a single reaction. Similar primer sets were reported in Babu et al. (2017) to produce intermediate length and full-length amplicons (57). In these experiments (Fig. 8F), all RNAs were detected in plants infected with RRV1-4 or RRV1-7. In this case, full-length amplicons corresponding to RNA2 were abundant. There were intermediate size amplicons of 1.8 and 2.0 kb, that are likely derivatives of RNA2 as reported in Babu et al. (2017). Notably, PCR products detecting RNA segments 3-7 co-migrate on a 1.2% agarose gel (Fig. 8F).

Reports indicate that the RRV produces double-strand RNA (dsRNA) in infected *Rosa multiflora* plants, and this has been used in the past to diagnose infection (58). Since the RT-PCR did not consistently detect the transcripts associated with the first four RNA segments in all samples, we carried out RNase protection assays using two samples of RRV1-4 infected plants (Fig. 8G). Total RNAs were treated with DNase I and then treated with 0, 1, and 5 units (U) of RNase I to digest single-strand RNA, leaving dsRNA intact. RT-PCR amplification using gene-specific primers was used to detect RNA segments 1 through 4 (Table 2). These PCR products ranged from 500 to 585 bp and confirm the presence of double-strand RNAs 1, 2, 3, and 4 are present in infected *N. benthamiana* plants but not in the mock-treated plants (Fig 8G). RNA from infected rose plants from garden rose samples produced similar RNAs 1-4 amplicons, while none were detected in healthy rose samples (Fig. 8G). These combined data indicate that the infectious clone is capable of producing systemic virus infection in *N. benthamiana* plants.

### Successful insertion of iLOV reporter into two locations in the genome to track systemic infection in whole plants

We fused the *iLOV* gene (335 bp; 10 kD protein) to the 3’ end of the Gc gene in RNA2 to serve as a fluorescent reporter to study virus accumulation, Gc functions and encapsidation (Fig. 9A). We replaced the wild-type RNA2 cDNA with the RNA2-iLOV in RRV1-4 (4R2-iLOV) and RRV1-7 (7R2-iLOV). Combining RRV1-4 with the RNA2-iLOV *N. benthamiana* plants were inoculated with 4R2-iLOV and 7R2-iLOV to determine if iLOV could be used to track infection. We monitored systemic movement of virus infection over 25 days and noted that the plants showed yellow mottling symptoms and wilting of the upper leaves, compared to the control (Fig. 9B). We monitored plants frequently using a hand-held UV lamp and failed to observe fluorescence in any leaves. However, the iLOV fluorescence was visible under epifluorescence microscopy at 13dpi, confirming systemic infection (Fig. 9B).

**FIG. 9.** *N. benthamiana* infected with 4R2-iLOV, 7R2-iLOV and 7R5-iLOV. A). Diagrammatic representation of iLOV fused to ORF2 on RNA2 and introduced into RNA5 replacing the ORF5. B). Images of plants at 25 dpi that are mock-inoculated or systemically infected with RRV 4R2-iLOV and 7R2-iLOV. Epifluorescence microscope images of infected leaves at 13 dpi shows fluorescence in epidermal cells. Bars= 100 µm. C). Images of plants at 25 and 35 dpi that are mock-inoculated or systemically infected with RRV 7R5-iLOV. Epifluorescence microscope images of infected leaves at 13 dpi shows fluorescence in epidermal cells. Bars= 100 µm. D). RT-PCR diagnostic using agRRV4-F1/R1 primer set to identify infected samples. PCR primers detecting EF1 as internal control showing equal loading in each lane. L=1 kb ladder. E). RT-PCR detecting each viral segment identified on the left. Samples infected with 4R2-iLOV, 7R2-iLOV and 7R5-iLOV are identified above the gels. EF-1 is a control PCR showing equal loading in each lane.

We hypothesized that RNA2 levels might not be as abundant as other segments, and this could account for the inability to view iLOV using a hand-held UV lamp. Therefore, we inoculated plants with all seven segments but used the R5-iLOV (whose ORF was replaced with iLOV cDNA but UTRs remain unchanged) in place of the wild-type R5 construct (Fig. 9A, C). We named this infectious clone 7R5-iLOV. Plants were infiltrated with the 7R5-iLOV, and systemic mottling and necrosis symptoms were evident at 25 dpi (Fig. 9C). Fluorescence was visualized at 13 dpi in the upper inoculated leaves using epifluorescence microscopy (Fig. 9D). Diagnostic RT-PCR using the agRRV4-F1/R1 primer set was performed to verify the presence of RRV in the upper leaves (Fig. 9D). These PCR products were sequenced to confirm the RRV sequences.

As with the prior experiments, we conducted additional RT-PCR to detect each genome segment confirming systemic infection at 25 dpi. Samples from the upper leaves of 4R2-iLOV, 7R2-iLOV, and 7R5-iLOV inoculated plants were analyzed by RT-PCR (Fig. 9E). In addition, we used the agRRV2-F1 and R2-iLOV-R primers (Table 4) to detect a 1080 bp band for RNA2 that overlaps the Gc-iLOV fusion. We used IF-agR5-F and iLOV-R primer pair to detect the iLOV sequences in RNA5, and this produced a 433 bp band (Fig. 9E). Both the recombinant RNA2 and RNA5 sequences containing iLOV were able to support infection (Fig. 9E).

### Successful confocal imaging of RRV Gc glycoprotein

The iLOV sequence was directly fused to 3’ end of the predicted RNA2 encoded glycoprotein precursor in the infectious clone (Fig. 10A). The predicted polyprotein is 73.78 kDa, and cleavage at the AKADD sequence would produce two mature 22.78 kD and 51 kDa glycoproteins known as the Gn and Gc proteins. By fusing the 10 kDa iLOV sequence to the C-terminus of the polyprotein, the mature Gc protein would be slightly extended. Since we lack antisera to iLOV or the Gc protein, we relied on confocal microscopy to characterize the chimeric Gc proteins. Plants were inoculated with 4R2-iLOV. At 7 dpi, the fluorescence in 4R2-iLOV infected cells was mainly cytosolic and nuclear. The large vacuole was void of fluorescence. There were fluorescent strands emanating from the nucleus and occasional spherical granules near these strands or along the periphery of the cell. These potentially represent virions or vesicles (Fig. 10B, C, D). We also inoculated plants with the R1_VAA_ construction, replacing the wild-type RNA1 in the 4R2-iLOV. At 7 dpi, we failed to detect iLOV fluorescence (Fig. 10E). The mock-inoculated leaves were agro-infiltrated to deliver the empty vector, and there was no iLOV fluorescence in these cells (Fig. 10F). The nuclear and cytoplasmic fluorescence was not expected. One explanation is that the polyprotein was further cleaved by cellular enzymes at the putative caspase3 cleavage site identified in Fig. 7.

**Fig. 10.** Visualization of Gc-iLOV in virus-infected cells by confocal microscopy. Scale bars represent 10 µm for all images. All confocal images are of epidermal cells at 7 dpi. A) Diagram represents the constructed cDNA with the iLOV fused to ORF2. The sizes of the mRNA and translated polyprotein is provided. After proteolysis at the conserved viral cleavage site motif the mature Gn and Gc-iLOV proteins are 22.75 and 61 kD. B, C, and D) Images of 4R2-iLOV infected epidermal cells. N identifies the nucleus. Arrows point to granular spheres. Red fluorescence is due to chloroplasts. E) Image of plants expressing 4R2-iLOV with the VAA mutation in the polymerase which suppresses replication. Cells show only red chloroplast fluorescence F). Image of mock-inoculated cells show only red chloroplast fluorescence. G-J) Images of 7R2-iLOV infected epidermal cells. Arrows point to granular bodies, and arrowheads identify fluorescence attached to chloroplasts. Images, J1, J2, J3, J4, are individual slices in a Z-stack that show the abundance of vesicles adjoining the nucleus and along a single transverse fluorescent strand that stretches across the cell.

Plants were also inoculated with 7R2-iLOV and imaged by confocal microscopy at 7 dpi. At low power, fluorescence was evident along the cell wall and in the nucleus and alongside strands stretching from the nucleus to the cell wall (Fig. 10G-J). Close examination of the chloroplasts revealed fluorescent halos around the chloroplast, suggesting that the Gc-iLOV fusion associates with the chloroplast membrane (Fig. 10G, H and I; arrowheads). The granular spheres that either represent vesicles or virions were more abundant in these cells. They seemed to gather near the nucleus, suggesting that they might bud from the nuclear envelope. Individual or aggregates of vesicles or virions were evident along fluorescent strands throughout the cell. These strands could be cytoskeletal, ER membranes, or cytoplasmic strands (Fig. 10 I, J1-J4). These first images of the viral glycoprotein suggest that it is dynamic in its association with various cellular structures and is influenced by the presence of other viral proteins.

### Inoculation using RRV infectious clone to rose plants

An infectious clone would be valuable to screen rose germplasm for resistance that can be incorporated into breeding programs. Since *Agrobacterium tumefaciens* causes crown gall disease in roses, we conducted a series of experiments to determine if agro-delivery of infectious RRV clones to plants produces RRV infection while eliminating the pathogenic effects of *A. tumefaciens*. For this, we compared the symptoms and timing of infection between plants inoculated with wild-type RRV contained in plant sap and agro-delivery of the RRV infectious clone. We used the infectious clone that did not contain the iLOV reporter to avoid the possibility of iLOV recombinants altering virus symptoms. A homogenate inoculum was prepared by grinding leaf tissue from naturally infected roses (cv. Single Knockout) in 0.05 M phosphate buffer plus 0.1% Silwett-77 and 1U/mL RNase Inhibitor (pH 7.0) (1:30 w/v). A set of healthy plants that had three canes were inoculated using an artist’s airbrush. Homogenate was delivered to compound leaves attached to the secondary stems of one cane (Fig. 11A). These secondary stems were surrounded by a shield to ensure that the airbrush delivery did not contaminate other leaves on other canes. This allowed us to sample non-inoculated leaves at a later time for evidence of virus spread through the plant. At the same time, we used the artist airbrush to agro-deliver RRV1-7 to leaves on the secondary stems of one cane using a similar shielding system. We monitored plants in the greenhouse for 60 days to understand the progression of symptoms and took photos at regular intervals. Pictures taken at 27 dpi showed mottled yellow leaves in virus-infected plants (Fig 11B, C, and D). These symptoms were consistent over the 60 days that the plants were observed. At the same time, we harvested one new emerging leaf from the inoculated cane and one leaf from the non-inoculated cane. We used the diagnostic RRV4F/R primer pair to verify that the plants were infected (Fig. 11E). All of the plants were systemically infected after inoculation with RRV containing plant sap or the infectious clones. We also took two samples from two plants that were treated with sap or the infectious clone and tested these with PCR primers to detect all seven segments. Only one plant that was treated with the infectious clone produced five of the seven PCR products. These data suggest that the RNA segments accumulate to different levels.

**Fig. 11.** Successful mechanical inoculation of RRV to roses. (A) Diagram representing airbrush method for delivery of plant homogenate or Agrobacterium cultures carrying infectious virus clone to a single cane, and the method for sampling new growth for infection. B, C, and D) Images show the mock-inoculated and virus-infected plants of ‘The Knock Out’® rose variety at 27 dpi. New growth shows yellow mosaic on the inoculated leaves. Veins remain dark green. E) RT-PCR verifies the presence of antigenomic RNA4 in two leaves on two different canes. The bars below the gel indicate the 2 samples from a single plant. S1 and S2 refer to PCR products from sap inoculated plants and 7R#1-7R#4 identify PCR products from plants inoculated with the infectious clone. L=molecular weight ladder. F) One rose plant that tested positive for RRV infection using the RRV1-7 infectious clone was used for PCR analysis to detect each RNA segment. RT-PCR failed to detect RNA1 and RNA2 (lanes 1 and 2), but detected RNA3, RNA4, RNA5, RNA6, and RNA7 (lanes 3-7 respectively) in inoculated rose plants. L= molecular weight ladder. G) Ultra-thin section of healthy rose leaf H) Images 1 and 2 show ultrathin section of garden infected rose leaf has a certain pathology. The internal symptoms appear as grey and dark grey spheres near the cell periphery and Golgi apparatus. These spheres have similar dimensions to virion particles but could be other pathological aggregates I, J) Aggregates of grey spherical structures in sap inoculated plants and infectious clone inoculated leaves, respectively Arrows in H-J point to examples of internal symptoms. The scale bars in each panel represent 0.5 *µ*m.

We conducted electron microscopy to look for evidence of internal symptoms or virions in RRV infected roses in the sap, and infectious clone treated plants. Internal symptoms could include abnormalities in organelles, changes in cell contents, or abnormal inclusions. Ultra-thin sections of resin embedded samples of infected garden rose leaves, sap inoculated rose leaves, and RRV1-7 infected leaves were prepared and stained with uranyl acetate and lead citrate. Ultra-thin sections of buffer-treated leaves were also prepared and stained. Images in Fig. 11G-J shows that the chloroplast, mitochondria, and cellular membranes are intact in control and infected samples. In all the infected garden rose samples, we observed electron-dense spheres near the cell periphery and adjacent structures that appear to be the Golgi apparatus. These spheres were either darkly stained or light grey and ranged from 0.8-1.0 µm in diameter. Because they were adjacent to Golgi and vesicles of similar dimensions, we could not discern virions from vesicles. These structures resemble the membranous structures reported in *Fig mosaic virus* (FMV) infected leaves by electron microscopy (59, 60) and these were seen only in virus-infected samples and not in tissues of healthy plants. Further experiments will require developing serological detection of the surface glycoprotein to validate the presence of virions in leaf tissues better. Overall, these data show that the infectious clone can be used to inoculate roses and produce systemic infection.

## DISCUSSION

For most *Bunyavirales* members, the RNA dependent RNA polymerase (RdRp) and nucleocapsid (N) proteins constitute the minimal protein machinery for genome replication and transcription. The N protein, which encapsidates the RNA genome to form the ribonucleoprotein (RNP) complex. The RNP is required for segment packaging during virus particle assembly. The N protein acts along with the viral RdRp for replication and mRNA synthesis(34, 61, 62). Also, the terminal untranslated regions (UTRs) are also critical for replication and transcription. The 5’ and 3’ UTRs are complementary and form a “panhandle” required for efficient protein synthesis in mini-replicon systems (26). Thus, identifying the RdRp and N proteins and characterizing the UTR regions was essential for building an RRV mini-replicon system as well as an infectious cDNA clone.

The phylogenies of the viral RdRp and N proteins point to an evolutionary relatedness with members of the *Orthobunyavirus genus*, such as BUNV and LACV, and to members of the *Hantavirus* genus, which are mostly arthropod or rodent borne viruses that cause significant diseases in humans and livestock (27, 29, 31). Surprisingly the plant infecting orthotospoviruses and tenuiviruses are more distant relatives. The alignment of amino acid sequences of the viral RdRps allowed us to identify the consensus sequences for the endonuclease domain near the N-terminus and the C-terminal polymerase motifs preA, A, B, C, D and E. The alignment of the amino acid sequences of the viral N proteins and the 3D models in combination revealed similarities to orthobunyaviruses with a central RNA binding cleft and extending N- and C-terminal arms that are likely involved in protein-protein interactions (32–34, 62, 63). We identified a number of charged Arg, His, and Lys residues in the central region that are likely to provide electrostatic bonds with the RNA backbone. These data provide an excellent framework for further reverse genetic studies to examine the role of the RRV N protein in replication and encapsidation(35). In the future we will develop antisera for immunological studies to conduct a broader range of experiments involving RNA binding, RNA synthesis, and genome encapsidation.

Further evidence of the relatedness of RRV to BUNV was found by examining the UTR regions of each genome segment. Each RRV segment has 13 conserved 3’ and 5’ nucleotides which are complementary. For BUNV the first 11 nucleotides are conserved and for RVFV it is the first 13 nucleotides that are conserved among segments (41, 42). In Fig. 4 we show that in all seven RRV segments the first eight nucleotides of the 3’ and 5’ ends are identical to BUNV. Nucleotide positions 10-13 on the 3’ UTR are also identical to BUNV. Studies of BUNV and other orthobunyaviruses determined that these sequences function as promoters for transcription and replication (64). These terminal UTR sequences contain signals for the viral polymerase to produce positive and negative sense replication as well as mRNA transcription. Studies of BUNV have shown that differences in the UTR regions also determine template activities for genome synthesis with some genome segments accumulating to higher levels in infected cells than others (42). If the UTRs in RRV control the accumulation of RNA segments in a similar manner to BUNV, then this could explain the variation in detection of various RNA segments by RT-PCR in infected *N. benthamiana* cells as seen in Figs 9 and 10. Detailed mutational analysis using a minireplicon system of BUNV-identified specific nucleotides within the 3’ and 5’ UTRs that control mRNA transcription (65). The mRNA is capped while genomic and antigenomic RNA is not capped. For orthobunyaviruses, the endonuclease activity of the viral RdRp cleaves a capped primer from host cell mRNAs (24). A study by Walia and Falk (2012) used 5’ RACE, RNA sequencing, and polyribosomal RNA extracted from FMV-infected leaves to show that viral mRNAs had 5’ heterogenous non-viral RNA sequences (59). The evidence of conserved UTR regions combined with cap-snatching mechanism for producing mRNAs further highlights the similarities between emaravirus and orthobunyaviruses.

The 5’ UTRs for RRV RNA2 through RNA7 were 190 to 631 nucleotides and each segment was determined to be monocistronic. RNA3, RNA4, and RNA6 have 5’ UTRs that are longer than the open reading frames. There is insufficient information yet to know if there is a possibility that the open reading frames could be extended by ribosome read-through of the stop codons. These genome segments are likely producing mRNAs with long 3’ UTRs making them very good targets for the cellular nonsense-mediated decay (NMD) pathway (66). This pathway normally acts to degrade aberrant mRNAs of the cell, but was recently reported to act on gRNA and subgenomic RNAs of viruses in the *Tombusviridae* family. Researchers showed that *Turnip crinkle virus* (TCV) gRNA and sgRNAs are subject to the NMD pathway because of their long UTRs (66). With the new RRV infectious clone and iLOV reporter technology, future research will be able to examine whether RRV transcripts are susceptible or resistant to NMD.

RRV belongs to the genus *Emaravirus* which is a recently described genus. *Rose rosette virus* encodes seven genome segments with negative sense polarity. Until now the tools have not existed to conduct functional analysis of the RRV proteins and their role in virus replication or plant-virus interactions. We prepared an agro-infectious clone of RRV that was able to systemically infect *N. benthamiana* and rose (cv Singleknock Out) plants. We introduced the iLOV reporter into RRV RNA5 as a template for studying the viral contributions to RNA synthesis. We demonstrated that this reporter is useful for monitoring virus accumulation in inoculated leaves either by microscopic observation or spectroscopic analysis. The iLOV which is a photostable flavin-based fluorescent protein that functions well in plants was reported to perform better than GFP as a reporter of plant virus infection when introduced into the *Tobacco mosaic virus* or *Pepino mosaic virus genomes* (12, 17). Its small size (335 bp coding region and 10kD protein), makes it an excellent choice to fuse to the glycoprotein and to replace the small open reading frame on RNA5 (467 bp) without significantly altering the size of the genome. We were concerned to use the *green fluorescent protein* (GFP), *luciferase*, or β*-glucuronidase* genes which are significantly larger sequences to insert into the genome. We considered that these other reporter genes might be less stable, might cause difficulties for genome packaging, or could alter virus symptoms in whole plants. Using the R5-iLOV construction, we demonstrated that the RdRp and N proteins were the minimal requirements to generate expression of the reporter protein from the recombinant RNA5 template. We also introduced a mutation into the putative RdRp replacing the conserved Ser-Asp-Asp tripeptide with Val-Ala-Ala tripeptide and this significantly hampered iLOV expression as seen using confocal microscopy and spectrophotometry. By fusing iLOV to the RNA2 encoded glycoprotein we were able to visualize Gc-iLOV associated structures by confocal microscopy. We also noted that the symptoms of virus infection were similar in plants inoculated with RRV sap, RRV infectious clone or the RRV infectious clone containing iLOV. However, fluorescence was barely detectable using a hand-held UV/blue lamp. These data suggest that further fluorescent protein alternatives need to be explored to improve the ability to monitor systemic spread of virus infection in whole plants. Overall, this research demonstrated that this reporter based reverse genetic system is suitable for studying overall RNA synthesis and subcellular associations of the viral glycoprotein.

The RNA2 polyprotein encodes the Gn and Gc proteins. Amino acid sequence analysis and 3D modeling predicts that the Gn protein has a signal peptide for membrane insertion and then two transmembrane domains (43, 44). These presumably drive insertion into the ER. The Gn also has a predicted ER export signal and Golgi retention signals in proximity to the predicted polyprotein cleavage site. The larger Gc protein has two potential ER export signals, as well as a single predicted C-terminal transmembrane domain and a single Golgi retention signal. These data suggest that the polyprotein may be inserted into the ER membranes and then cleaved into two proteins which remain interacting. The Gn and Gc complex move are exported from the ER and retained in the Golgi apparatus. It was interesting that we detected spherical structures that were similar in dimension to virions by electron microscopy that were close to Golgi and other vesicles which lends support to the endomembrane relationships predicted by the alignments and the 3D model. Fusing iLOV to the Gc protein enabled us to view small spherical bodies that might be vesicles or virions in *N. benthamiana* leaves which often appeared along strands extending from the nucleus to the cell periphery. The success of establishing a fluorescent tag to follow RRV infection opens up new possibilities to study virion maturation and the formation of enveloped particles. The structures seen in this study by electron microscopy and confocal microscopy resemble structures associated with the FMV nucleoprotein complex that was previously reported. This alternative RRV infectious clone will be used for mutational analysis to characterize signal motifs affecting subcellular accumulation and proteolytic maturation of the Gc protein.

Another approach to tagging virion glycoproteins was conducted using BUNV (67). Here the enhanced GFP or mCherry fluorescent protein sequences were inserted into the glycoprotein precursor replacing the N-terminal fragment of Gc. The chimeric Gc was useful for visualizing BUNV cellular attachment, assembly and budding to track intracellular trafficking. The use of iLOV as a C-terminal tag allows for similar research without having to truncate the protein to achieve the chimera, however it is worth considering locating iLOV at the N-terminus of Gc to see if an alternative location will provide different insights into virion assembly and intracellular trafficking.

The majority of the literature relating to members of the genus *Emaravrius* either report new species or new detection methods. There are only a few reports that functionally analyzed emaravirus proteins by overexpressing them from heterologous plasmids or viruses which identified the N protein of FMV, the movement protein of *Raspberry leaf blotch virus*, and two silencing suppressor proteins of *High plains wheat mosaic virus* P7 and P8 (56, 60, 68). Until now a reverse genetic approach and live imaging of these viruses was not possible. We demonstrated here that RRV replication machinery is quite similar to BUNV and other orthobunyaviruses. Given these similarities we hope to use RRV as a plant virus model for discovery of conserved eukaryotic host factors that contribute to virus replication or defend against infection. The ability to fluorescently tag RRV by introducing iLOV into two genomic locations shows that this virus is amenable to manipulations that will allow us to explore various aspects of infection and provides expanding opportunities to visualize the primary infection sites in the inoculated leaves. We found that RRV infection is difficult to detect by molecular methods other than RT-PCR in inoculated and systemic leaves, which may be caused by already low viral RNA levels. We also found that iLOV was not bright enough to the naked eye for use as a visual marker to track systemic infection from leaf to leaf. While the results of this study raise further questions about the mechanisms for RNA synthesis and virion assembly, we provide the first tools to begin to address these new questions. As we expand our resources and tools for further investigations, the replicon system and infectious clone will be valuable for dissection of RRV protein functions.

## MATERIALS AND METHODS

### Bioinformatic analyses

Protein sequences of viruses were retrieved from the National Center for Biotechnology Information (NCBI) protein archive (https://www.ncbi.nlm.nih.gov/protein). Sequences quality was manually checked using Geneious Prime® 2019.2.1, and multiple sequence alignment was carried out using Muscle ver 3.8.425 (maximum number of iterations at 8) build into the Genious Prime software (https://www.geneious.com) (69). All phylogenetic trees were generated using model VT+F+G4 with 1000 ultrafast bootstraps using IQ-TREE ver. 1.6.11 (28). Phylogenetic trees were visualized using iTOL ver 4.4.2 (70). Protein domain analysis for the RdRp was carried out by aligning a subset of protein using CLC Workbench and examining reported alignments and crystallographic structures of well-studied *Bunyavirales*. The identified highly conserved sequences are annotated in Fig. 2 (29–31). The N protein folded structures were generated and analyzed using I-Tasser (38) and The PyMOL Molecular Graphics System 2.0 (Schrodinger, LLC). The list of species and their Genbank Accession numbers is provided in Table S2. The mFOLD v3.6 Web Server (40)was used to characterize the complementary 3’ and 5’ UTRs and provide the free energy determination. The glycoprotein precursor folded structures were generated and analyzed using I-Tasser and Pro-Origami (38, 48). The list of species and their GenBank accession numbers is provided in Table S1.

### Construction of RRV infectious clones

To prepare the RRV infectious clone, the full-length cDNAs for agRNA1 (7026 bp), agRNA2 (2245 bp), agRNA3 (1544 bp) and agRNA4 (1541 bp) were synthesized *de novo* (NCBI reference sequences NC_015298.1, NC_015299.1, NC_015300.1, and NC_015301.1) and inserted into the small binary plasmid pCB301-HDV, which contains the double CaMV 35S promoter, HDRz, and Nos terminator (18) by GenScript (Piscataway, NJ). Plasmids were named pCB301-agRNA1, -agRNA2, -agRNA3, and -agRNA4. The pCB301-HDV plasmid is a binary plasmid with a duplicated cauliflower mosaic virus (CaMV) 35S promoter and 3’ HDRz sequence which was provided by Dr. Zhenghe Li (Zhejiang University, Hangzhou China). The full-length agRNA segments 5, 6 and 7 (MN095111, MN095112, MN095113) were amplified by RT-PCR prepared from infected rose leaves (var ‘Julia Child’) using the following primer pairs: IF_agR5F/R, IF_agR6F/R, and IF_agR7F/R with 15 nt that overlap vector sequences (Table 4). The antigenomic cDNA was positioned next to the CaMV 35S promoter and HDRz to produce viral transcripts with authentic 5’ and 3’ ends. All constructs were confirmed by sequencing before being transformed into *A. tumefaciens*.

To prepare pCB301_R2 iLOV and pCB301_R5 iLOV constructs, the primers of R2_iLOV.F/R with pCB301_R2.F/R and R5_iLOV.F/R with pCB301_R5.F/R (Table 1) were used to amplify iLOV fragments from TMV_iLOV (17) and fragments of pCB301_R2 and pCB301_R5. All PCRs were carried out using high-fidelity 2X Platinum SuperFi® Green PCR Master Mix (Invitrogen). The high fidelity directional In-Fusion® HD Cloning Kit (Takara Bio USA, Inc.) was used to introduce each amplified full-length cDNA into the pCB301-HDV.

### Plant materials and virus inoculation

*Nicotiana benthamiana* were grown at 23°C with 16 h/ 8 h (day/night) photoperiod in a growth chamber. Rose plants (cvs. Knockout White and Knockout Pink) were grown in a greenhouse with temperature settings to at 23°C. Three-weeks-old plants were inoculated with extracts taken from rose rosette virus-infected rose plants (var. Julia Child). Plant extracts were prepared by grinding 0.5 g infected leaves in 15 mL (1:30 w/v) 0.05 M phosphate buffer (pH 7.0) and filtered through cheesecloth. Then SUPERaseIn(tm) RNase inhibitor (Invitrogen) (1 U/ml) and 0.1% volume of Silwet-77 were added to the extract and loaded to the reservoir of an artist airbrush. Plants were lightly dusted with carborundum, and extracted leaf homogenate (from Single Knockout roses) was applied using the Central Pneumatic® 3/4 a 3 oz airbrush kit (Harbor Freight, Plano, TX) (Fig. 1A).

Three-weeks-old plants were also infiltrated with *A. tumefaciens* cultures harboring constructs for each RRV agRNA segment. Cultures were grown overnight in YEP media, resuspended in MES buffer (10 mM MgCl_2_, 10 mM MES, pH 5.6, and 150 uM acetosyringone), and adjusted to an optical density A_600_ of 1.0. After 2-4 hours of incubation in the dark, equal volumes of each *Agrobacterium* culture were mixed at 1.0 OD and loaded to a 1-mL syringe for infiltration to *N. benthamiana*. Mixed *Agrobacterium* cultures were delivered to rose plants using the Central Pneumatic® airbrush.

### RNA isolation, RT-PCR, and RNaseI protection assays

Total RNA was extracted from leaves using the Maxwell® 16 Instrument LEV simplyRNA Purification Kits (Promega). All reverse transcription reactions were carried out using random primers and the High-capacity cDNA reverse transcription kit (Applied Biosystems®). PCR amplification was performed using GoTaq® G2 master mixes DNA polymerase (Promega) with the following primer pairs: agRRV1-F1/R1, agRRV2-F1/R1, agRRV3-F1/R1, agRRV3-F2/R2, agRRV4-F1/R1, agRRV5-F1/R1, agRRV5-F1/R1, agRRV6-F1/R1 and agRRV7-F1/R1 (Table 1). We used primer pairs to detect EF1 transcripts in *N. benthamiana* and Actin in roses as internal controls for RT-PCR verification of virus infected plants (Table 1). RNase I protection assays were performed using total RNA (5 µg) from *N. benthamiana* leaves, as well as infected or healthy garden rose samples according to (71). Samples were treated with DNase I for 1 h at 37°C. RNase I-treated RNA was reverse transcribed using random primers and then amplified using the following primer pairs: agRRV1-F1/R1, agRRV2-F1/R1, agRRV3-F2/R2, and agRRV4-F1/R1 (Table 4). All PCR amplified products were subjected to 1.2% agarose gel electrophoresis stained with ethidium bromide (71).

### Epifluorescence, confocal, and transmission electron microscopy

Leaf segments were cut with a scalpel and places on a microscope slide and coverslip and then examined with a Nikon Eclipse 90i epifluorescence microscope using the FITC (Fluorescein Isothiocyanate) filter (excitation 465nm/495nm; emission 515nm/555nm) to detect YFP, GFP, and iLOV. Images were captured using a DS-Ri1 camera and NIS-Elements AR-3.2 software (Nikon). An Olympus FV1000 confocal microscope was used for high-resolution imaging of leaf segments.

Rose leaf samples were resin embedded, and ultrathin sections were prepared by the Texas A&M University Microscopy and Imaging Core Facility. Where indicated, the Pelco Biowave microwave processor (Ted Pella, Inc., Redding, CA) equipped with ColdSpot® technology was used, with temperature set to 20 °C. Microwave power settings are expressed as continuous power in watts (W). All fixation and washing steps, as well as epoxy resin infiltration with microwave processing, were performed under vacuum, while the dehydration steps were performed without vacuum. Leaves were cut into 1 mm square sections, immersed in Trump’s fixative in a polypropylene microcentrifuge tube and vacuum infiltrated to remove air from the tissue (72). Samples were kept at room temperature for 30 minutes, then microwaved at 150 W with a 1-3-1-2 cycle (magnetron 1 min on, 3 min off, 1 min on, 2 min off) under vacuum and then microwaved at 650 W with a 10s on, 20s off, 10 s on cycle (73). Samples were washed three times in Trump’s fixative and twice in water for 1 min at 150W each time, then post-fixed in 1% aqueous osmium tetroxide at 100W power, using five 2 min on −2 min off cycles. Osmium tetroxide was then replaced by 2.5% w/v aqueous potassium ferrocyanide (Sigma-Aldrich), and five 2-2 cycles were performed as above. This sequential osmium post-fixation and reduction treatment yielded better contrast without overstaining (74), compared to the simultaneous treatment with reduced osmium. Samples were then washed in water twice for 1 min at 250 W, en-bloc stained with 2% w/v aqueous uranyl acetate a 100 W, using three 2 min on, 2 min off cycles, and rinsed twice in water as above. Dehydration in acetone series (10, 20, 40, 60, 80% v/v acetone in water) and two times in 100% acetone was performed for 3 min at 200 W, followed by 10 minutes on a rotator in each step. Infiltration with modified Quetol/Spurr’s low viscosity resin (75) was performed as follows: Acetone: resin 1:1 mixture, 3 min at 200W without vacuum; acetone: resin 1:1 mixture, acetone: resin 1:2 mixture, 3 min at 200 W under vacuum each step. This was followed by four changes of 100% resin. Each time, samples were kept on a rotator for 1-2 h, then microwaved under vacuum for 5 min at 200W. Samples were then transferred to embedding molds and polymerized at 60 °C for 2 days.

Ultrathin sections were collected on 200-mesh copper grids, stained with 2% uranyl acetate for 5 min, and freshly prepared Reynold’s lead citrate 5 min. Images were acquired on a JEOL 1200Ex transmission electron microscope at calibrated magnification, using SIA 15C CCD camera

### Fluorometry

*N. benthamiana* leaves were agro-infiltrated with three combinations of pCB301 constructs: RNA1, RNA3, and RNA5-iLov; RNA1_VAA_, RNA3, and RNA5-iLov; or mock (empty pCB301). Fluorometry was used to assess the expression level of the fluorescent protein iLOV in the three biological replicates. At 3 dpi, 40 mg of infiltrated tissue was harvested into microtubes and ice cold 0.05M phosphate-buffered saline (PBS) was added to make a 1:5 (w:v) dilution. The samples were homogenized with a micropestle, then centrifuged at 6k rpm for 8 minutes. Then 20uL of the supernatant were loaded into AppliedBiosystems’ MicroAmp EnduraPlate Optical 96-Well Fast Clear Reaction Plate, and the samples were analyzed using the spectroscopic capabilities of the AppliedBiosystems’ QuantStudio 3 Real-Time PCR System. Fluorescence was determined by collecting emission wave amplitude after heating the sample to 25C for 15 seconds. The filter set used was for SYBR Green I reporter (497/520 nm excitation/emission). Raw data were normalized to the fluorescence level of a healthy leaf sample and mock treatments using Microsoft Excel. The Pierce Coomassie (Bradford) protein assay reagent (ThermoFisher) was used to measure the total protein concentration of each sample. Albumin standards were used to generate a standard curve for determining the concentrations of each unknown sample. The Fluorescence Units (FU) per μg protein were reported.

## ACKNOWLEDGMENTS

The authors declare no conflict of interest. We thank the Rose Rosette Disease Network for their support for this research. We thank Dr. Li Zhenghe from Zhejiang University for providing the pCB301-HDV plasmid for our constructions. Michele Scheiber at Star Roses and Plants for providing rose liners for experiments. The American Rose Society for providing financial gifts that enabled this research. USDA’s National Institute of Food and Agriculture (NIFA) Specialty Crop Research Initiative project “Combating Rose Rosette Disease: Short Term and Long-Term Approaches” (2014-51181-22644/ SCRI). We also thank Stanislav Vitha and Rick Littleton of the Texas A&M Microscopy and Imaging Center for assistance with electron and confocal microscopy. Stanislav Vitha developed the methods for resin embedding rose tissues and Rick Littleton provided the ultrathin sections, staining, and training.

## REFERENCES

1. Ahlquist P, French R, Janda M, Loesch-Fries LS. 1984. Multicomponent RNA plant virus infection derived from cloned viral cDNA. Proc Natl Acad Sci U S A 81:7066–7070.

2. Desbiez C, Chandeysson C, Lecoq H, Moury B. 2012. A simple, rapid and efficient way to obtain infectious clones of potyviruses. J Virol Methods.

3. Bedoya LC, Daròs JA. 2010. Stability of Tobacco etch virus infectious clones in plasmid vectors. Virus Res 149:234–240.

4. Flatken S, Ungewickell V, Menzel W, Maiss E. 2008. Construction of an infectious full-length cDNA clone of potato virus M. Arch Virol 153:1385–1389.

5. Ali MC, Omar AS, Natsuaki T. 2011. An infectious full-length cDNA clone of potato virus Y NTN-NW, a recently reported strain of PVY that causes potato tuber necrotic ringspot disease. Arch Virol 156:2039–2043.

6. Bordat A, Houvenaghel MC, German-Retana S. 2015. Gibson assembly: An easy way to clone potyviral full-length infectious cDNA clones expressing an ectopic VPg Plant viruses. Virol J 12:1–8.

7. Lindbo JA. 2007. TRBO: A high-efficiency Tobacco mosaic virus RNA-based overexpression vector. Plant Physiol 145:1232–1240.

8. Boyer J-C, Haenni A-L. 1994. Infectious transcripts and cDNA clones of RNA viruses. Virology 198:415–426.

9. Dolja V V., McBride HJ, Carrington JC. 1992. Tagging of plant potyvirus replication and movement by insertion of β-glucuronidase into the viral polyprotein. Proc Natl Acad Sci U S A 89:10208–10212.

10. Mardanova ES, Kotlyarov RY, Kuprianov V V., Stepanova LA, Tsybalova LM, Lomonosoff GP, Ravin N V. 2015. Rapid high-yield expression of a candidate influenza vaccine basedon the ectodomain of M2 protein linked to flagellin in plants using viral vectors. BMC Biotechnol 15:1–10.

11. Tian J, Pei H, Zhang S, Chen J, Chen W, Yang R, Meng Y, You J, Gao J, Ma N. 2014. TRV-GFP: A modified Tobacco rattle virus vector for efficient and visualizable analysis of gene function. J Exp Bot 65:311–322.

12. Sempere RN, Gómez P, Truniger V, Aranda MA. 2011. Development of expression vectors based on pepino mosaic virus. Plant Methods 7:6.

13. Stevens M, Viganó F. 2007. Production of a full-length infectious GFP-tagged cDNA clone of Beet mild yellowing virus for the study of plant-polerovirus interactions. Virus Genes 34:215–221.

14. Rabindran S, Dawson WO. 2001. Assessment of recombinants that arise from the use of a TMV-based transient expression vector. Virology 284:182–189.

15. Zhao Y, Hammond J, Tousignant ME, Hammond RW. 2000. Development and evaluation of a complementation-dependent gene delivery system based on cucumber mosaic virus. Arch Virol 145:2285–2295.

16. Shivprasad S, Pogue GP, Lewandowski DJ, Hidalgo J, Donson J, Grill LK, Dawson WO. 1999. Heterologous sequences greatly affect foreign gene expression in tobacco mosaic virus-based vectors. Virology 255:312–323.

17. Chapman S, Faulkner C, Kaiserli E, Garcia-Mata C, Savenkov EI, Roberts AG, Oparka KJ, Christie JM. 2008. The photoreversible fluorescent protein iLOV outperforms GFP as a reporter of plant virus infection. Proc Natl Acad Sci 105:20038–20043.

18. Wang Q, Ma X, Qian S, Zhou X, Sun K, Chen X, Zhou X, Jackson AO, Li Z. 2015. Rescue of a plant negative-strand RNA virus from cloned cDNA: Insights into enveloped plant virus movement and morphogenesis. PLOS Pathog 11:e1005223.

19. Ganesan U, Bragg JN, Deng M, Marr S, Lee MY, Qian S, Shi M, Kappel J, Peters C, Lee Y, Goodin MM, Dietzgen RG, Li Z, Jackson AO. 2013. Construction of a Sonchus yellow net virus minireplicon: A step toward reverse genetic analysis of plant negative-strand RNA viruses. J Virol 87:13081–13081.

20. Schnell MJ, Mebatsion T, Conzelmann KK. 2018. Infectious rabies viruses from cloned cDNA. EMBO J 13:4195–4203.

21. Jackson AO, Dietzgen RG, Goodin MM, Li Z. 2018. Development of model systems for plant Rhabdovirus researchAdvances in Virus Research, 1st ed. Elsevier Inc.

22. Abudurexiti A, Adkins S, Alioto D, Alkhovsky S V., Avšič-Županc T, Ballinger MJ, Bente DA, Beer M, Bergeron É, Blair CD, Briese T, Buchmeier MJ, Burt FJ, Calisher CH, Cháng C, Charrel RN, Choi IR, Clegg JCS, de la Torre JC, de Lamballerie X, Dèng F, Di Serio F, Digiaro M, Drebot MA, Duàn X, Ebihara H, Elbeaino T, Ergünay K, Fulhorst CF, Garrison AR, Gāo GF, Gonzalez J-PJ, Groschup MH, Günther S, Haenni A-L, Hall RA, Hepojoki J, Hewson R, Hú Z, Hughes HR, Jonson MG, Junglen S, Klempa B, Klingström J, Kòu C, Laenen L, Lambert AJ, Langevin SA, Liu D, Lukashevich IS, Luò T, Lǚ C, Maes P, de Souza WM, Marklewitz M, Martelli GP, Matsuno K, Mielke-Ehret N, Minutolo M, Mirazimi A, Moming A, Mühlbach H-P, Naidu R, Navarro B, Nunes MRT, Palacios G, Papa A, Pauvolid-Corrêa A, Pawęska JT, Qiáo J, Radoshitzky SR, Resende RO, Romanowski V, Sall AA, Salvato MS, Sasaya T, Shěn S, Shí X, Shirako Y, Simmonds P, Sironi M, Song J-W, Spengler JR, Stenglein MD, Sū Z, Sūn S, Táng S, Turina M, Wáng B, Wáng C, Wáng H, Wáng J, Wèi T, Whitfield AE, Zerbini FM, Zhāng J, Zhāng L, Zhāng Y, Zhang Y-Z, Zhāng Y, Zhou X, Zhū L, Kuhn JH. 2019. Taxonomy of the order Bunyavirales: update 2019. Arch Virol 164:1949–1965.

23. Mielke-Ehret N, Mühlbach HP. 2012. Emaravirus: Anovel genus of multipartite, negative strand RNA plant viruses. Viruses.

24. Sun Y, Li J, Gao GF, Tien P, Liu W. 2018. Bunyavirales ribonucleoproteins: the viral replication and transcription machinery. Crit Rev Microbiol 44:522–540.

25. Waliczek TM, Byrne D, Holeman D. 2018. Opinions of landscape roses available for purchase and preferences for the future market. Horttechnology 28:807–814.

26. Pemberton HB, Ong K, Windham M, Olson J, Byrne DH. 2018. What is rose rosette disease? HortScience 53:592–595.

27. Klemm C, Reguera J, Cusack S, Zielecki F, Kochs G, Weber F. 2013. Systems to establish Bunyavirus genome replication in the absence of transcription. J Virol 87:8205–8212.

28. Trifinopoulos J, Nguyen LT, von Haeseler A, Minh BQ. 2016. W-IQ-TREE: a fast online phylogenetic tool for maximum likelihood analysis. Nucleic Acids Res.

29. Amroun A, Priet S, de Lamballerie X, Quérat G. 2017. Bunyaviridae RdRps: structure, motifs, and RNA synthesis machinery. Crit Rev Microbiol 43:753–778.

30. Kormelink R, Garcia ML, Goodin M, Sasaya T, Haenni AL. 2011. Negative-strand RNA viruses: The plant-infecting counterparts. Virus Res 162:184–202.

31. Gerlach P, Malet H, Cusack S, Reguera J. 2015. Structural insights into bunyavirus replication and its regulation by the vRNA promoter. Cell 161:1267–1279.

32. Guo Y, Wang W, Sun Y, Ma C, Wang X, Wang X, Liu P, Shen S, Li B, Lin J, Deng F, Wang H, Lou Z. 2016. Crystal structure of the core region of Hantavirus nucleocapsid protein reveals the mechanism for ribonucleoprotein complex formation. J Virol 90:1048–1061.

33. Xu X, Severson W, Villegas N, Schmaljohn CS, Jonsson CB. 2002. The RNA binding domain of the Hantaan virus N protein maps to a central, conserved region. J Virol 76:3301–3308.

34. Guo Y, Liu B, Ding Z, Li G, Liu M, Zhu D, Sun Y, Dong S, Lou Z. 2017. Distinct mechanism for the formation of the ribonucleoprotein complex of Tomato spotted wilt virus. J Virol 91:1–14.

35. Ariza A, Tanner SJ, Walter CT, Dent KC, Shepherd DA, Wu W, Matthews S V., Hiscox JA, Green TJ, Luo M, Elliott RM, Fooks AR, Ashcroft AE, Stonehouse NJ, Ranson NA, Barr JN, Edwards TA. 2013. Nucleocapsid protein structures from orthobunyaviruses reveal insight into ribonucleoprotein architecture and RNA polymerization. Nucleic Acids Res 41:5912–5926.

36. Komoda K, Narita M, Yamashita K, Tanaka I, Yao M. 2017. Asymmetric trimeric ring structure of the nucleocapsid protein of Tospovirus. J Virol 91:1–13.

37. Olaya C, Adhikari B, Raikhy G, Cheng J, Pappu HR. 2019. Identification and localization of Tospovirus genus-wide conserved residues in 3D models of the nucleocapsid and the silencing suppressor proteins. Virol J 16:1–15.

38. Yang J, Zhang Y. 2015. I-TASSER server: new development for protein structure and function predictions. Nucleic Acids Res 43:W174–W181.

39. Yang J, Zhang Y. 2015. Protein structure and function prediction using I-TASSER. Curr Protoc Bioinforma.

40. Zuker M. 2003. Mfold web server for nucleic acid folding and hybridization prediction. Nucleic Acids Res.

41. Barr JN, Elliott RM, Dunn EF, Wertz GW. 2003. Segment-specific terminal sequences of Bunyamwera bunyavirus regulate genome replication. Virology 311:326–338.

42. Barr JN, Rodgers JW, Wertz GW. 2005. The Bunyamwera virus mRNA transcription signal resides within both the 3’ and the 5’ terminal regions and allows ambisense transcription from a model RNA segment. J Virol 79:12602–12607.

43. Guardado-Calvo P, Rey FA. 2017. The Envelope Proteins of the Bunyavirales. Adv Virus Res 98:83–118.

44. Halldorsson S, Behrens AJ, Harlos K, Huiskonen JT, Elliott RM, Crispin M, Brennan B, Bowden TA. 2016. Structure of a phleboviral envelope glycoprotein reveals a consolidated model of membrane fusion. Proc Natl Acad Sci U S A 113:7154–7159.

45. Albornoz A, Hoffmann A, Lozach P-Y, Tischler N. 2016. Early Bunyavirus-host cell interactions. Viruses 8:143.

46. Banerjee N, Mukhopadhyay S. 2016. Viral glycoproteins: biological role and application in diagnosis. VirusDisease 27:1–11.

47. Gerrard SR, Nichol ST. 2002. Characterization of the Golgi retention motif of Rift valley fever virus Gn glycoprotein. J Virol 76:12200–12210.

48. Stivala A, Wybrow M, Wirth A, Whisstock JC, Stuckey PJ. 2011. Automatic generation of protein structure cartoons with pro-origami. Bioinformatics.

49. Guardado-Calvo P, Rey FA. 2017. The envelope proteins of the Bunyavirales. Adv Virus Res 98:83–118.

50. McGavin WJ, Mitchell C, Cock PJA, Wright KM, MacFarlane SA. 2012. Raspberry leaf blotch virus, a putative new member of the genus Emaravirus, encodes a novel genomic RNA. J Gen Virol 93.

51. Kumar S, Subbarao B, Hallan V. 2017. Molecular characterization of emaraviruses associated with Pigeonpea sterility mosaic disease. Sci Rep 7:1–20.

52. Patil BL, Kumar PL. 2015. Pigeonpea sterility mosaic virus: A legume-infecting Emaravirus from South Asia. Mol Plant Pathol 16:775–786.

53. Xin M, Cao M, Liu W, Ren Y, Zhou X, Wang X. 2017. Two negative-strand RNA viruses identified in watermelon represent a novel clade in the order Bunyavirales. Front Microbiol 8:1514.

54. Cifuentes-Muñoz N, Salazar-Quiroz N, Tischler ND. 2014. Hantavirus Gn and Gc envelope glycoproteins: Key structural units for virus cell entry and virus assembly. Viruses 6:1801–1822.

55. Dessau M, Modis Y. 2013. Crystal structure of glycoprotein C from Rift Valley fever virus. Proc Natl Acad Sci U S A 110:1696–1701.

56. Yu C, Karlin DG, Lu Y, Wright K, Chen J, MacFarlane S. 2013. Experimental and bioinformatic evidence that raspberry leaf blotch emaravirus P4 is a movement protein of the 30K superfamily. J Gen Virol 94:2117–2128.

57. Babu B, Washburn BK, Ertek TS, Miller SH, Riddle CB, Knox GW, Ochoa-Corona FM, Olson J, Katircioğlu YZ, Paret ML. 2017. A field based detection method for Rose rosette virus using isothermal probe-based Reverse transcription-recombinase polymerase amplification assay. J Virol Methods 247:81–90.

58. Di, R., Hill, J. H., and Epstein AH. 1990. Double-stranded RNA associated with the rose rosette disase of multiflora rose.

59. Walia JJ, Falk BW. 2012. Fig mosaic virus mRNAs show generation by cap-snatching. Virology 426:162–166.

60. Ishikawa K, Miura C, Maejima K, Komatsu K, Hashimoto M, Tomomitsu T, Fukuoka M, Yusa A, Yamaji Y, Namba S. 2015. Nucleocapsid protein from Fig mosaic virus forms cytoplasmic agglomerates that are hauled by endoplasmic reticulum streaming. J Virol 89:480–491.

61. Pyle JD, Whelan SPJ. 2019. RNA ligands activate the Machupo virus polymerase and guide promoter usage. Proc Natl Acad Sci U S A 116:10518–10524.

62. Wichgers Schreur PJ, Kormelink R, Kortekaas J. 2018. Genome packaging of the Bunyavirales. Curr Opin Virol 33:151–155.

63. Walter CT, Costa Bento DF, Guerrero Alonso A, Barr JN. 2011. Amino acid changes within the Bunyamwera virus nucleocapsid protein differentially affect the mRNA transcription and RNA replication activities of assembled ribonucleoprotein templates. J Gen Virol 92:80–84.

64. Ferron F, Weber F, de la Torre JC, Reguera J. 2017. Transcription and replication mechanisms of Bunyaviridae and Arenaviridae L proteins. Virus Res 234:118–134.

65. Weber F, Dunn EF, Bridgen A, Elliott RM. 2001. The Bunyamwera virus nonstructural protein NSs inhibits viral RNA synthesis in a minireplicon system. Virology 281:67–74.

66. May JP, Yuan X, Sawicki E, Simon AE. 2018. RNA virus evasion of nonsense-mediated decay. PLoS Pathog 14:1–22.

67. Shi X, van Mierlo JT, French A, Elliott RM. 2010. Visualizing the replication cycle of Bunyamwera Orthobunyavirus expressing fluorescent protein-tagged Gc glycoprotein. J Virol 84:8460–8469.

68. Gupta AK, Hein GL, Tatineni S. 2019. P7 and P8 proteins of High plains wheat mosaic virus, a negative-strand RNA virus, employ distinct mechanisms of RNA silencing suppression. Virology 535:20–31.

69. Edgar RC. 2004. MUSCLE: Multiple sequence alignment with high accuracy and high throughput. Nucleic Acids Res 32:1792–1797.

70. Letunic I, Bork P. 2019. Interactive Tree Of Life (iTOL) v4: recent updates and new developments. Nucleic Acids Res 47:W256–W259.

71. Okano Y, Senshu H, Hashimoto M, Neriya Y, Netsu O, Minato N, Yoshida T, Maejima K, Oshima K, Komatsu K, Yamaji Y, Namba S. 2014. In planta recognition of a double-stranded RNA synthesis protein complex by a potexviral RNA silencing suppressor. Plant Cell 26:2168–2183.

72. McDowell EM, Trump BF. 1976. Histologic fixatives suitable for diagnostic light and electron microscopy. Arch Pathol Lab Med.

73. Ferris AM, Giberson RT, Sanders MA, Day JR. 2009. Advanced laboratory techniques for sample processing and immunolabeling using microwave radiation. J Neurosci Methods.

74. Hua Y, Laserstein P, Helmstaedter M. 2015. Large-volume en-bloc staining for electron microscopy-based connectomics. Nat Commun.

75. Ann Ellis E. 2006. Solutions to the problem of substitution of ERL 4221 for vinyl cyclohexene dioxide in Spurr low viscosity embedding formulations. Micros Today.

76. Kalyaanamoorthy S, Minh BQ, Wong TKF, Von Haeseler A, Jermiin LS. 2017. ModelFinder: Fast model selection for accurate phylogenetic estimates. Nat Methods 14:587–589.

77. Nguyen LT, Schmidt HA, Von Haeseler A, Minh BQ. 2015. IQ-TREE: A fast and effective stochastic algorithm for estimating maximum-likelihood phylogenies. Mol Biol Evol 32:268–274.

78. Hoang DT, Chernomor O, Von Haeseler A, Minh BQ, Vinh LS. 2018. UFBoot2: Improving the ultrafast bootstrap approximation. Mol Biol Evol 35:518–522.

79. Babu B, Washburn BK, Poduch K, Knox GW, Paret ML. 2016. Identification and characterization of two novel genomic RNA segments RNA5 and RNA6 in rose rosette virus infecting roses. Acta Virol 60:156–165.

